# CD26-negative and CD26-positive tissue-resident fibroblasts contribute to functionally distinct CAF subpopulations in breast cancer

**DOI:** 10.1101/2022.09.27.509697

**Authors:** Julia M. Houthuijzen, Roebi de Bruijn, Eline van der Burg, Anne Paulien Drenth, Ellen Wientjens, Tamara Filipovic, Esme Bullock, Chiara S. Brambillasca, Marja Nieuwland, Iris de Rink, Frank van Diepen, Sjoerd Klarenbeek, Ron Kerkhoven, Valerie G. Brunton, Colinda L.G.J. Scheele, Mirjam C. Boelens, Jos Jonkers

## Abstract

Cancer-associated fibroblasts (CAFs) are abundantly present in the microenvironment of virtually all tumors and strongly impact tumor progression. Despite increasing insight into their function and heterogeneity, little is known regarding the origin of CAFs. Understanding the origin of CAF heterogeneity is needed to develop successful CAF-based targeted therapies. Through various transplantation studies in mice we determined that CAFs in both invasive lobular breast cancer and triple negative breast cancer originate from mammary tissue-resident normal fibroblasts (NFs). Single-cell transcriptomics, *in vivo* tracing and *in vitro* studies revealed the transition of CD26+ and CD26- NF populations into inflammatory CAFs (iCAFs) and myofibroblastic CAFs (myCAFs), respectively. *In vitro* functional assays showed that CD26+ NFs transition into pro-tumorigenic iCAFs which recruit myeloid cells in a CXCL12-dependent manner and enhance tumor cell invasion via matrix-metalloproteinase (MMP) activity. Together, our data show that CD26+ and CD26- NFs transform into distinct CAF subpopulations in breast cancer.

## Introduction

Tumorigenesis is not only governed by genetically altered cancer cells *per se*, but also by non-malignant host cells in the tumor microenvironment (TME), which strongly impact tumor progression, metastasis, and therapy response[1]. One of the most dominant cell types within the TME are the cancer-associated fibroblasts (CAFs). Decades of research has shown that CAFs can affect all hallmarks of cancer[2]. However, to label CAFs as the ‘bad guys’ within the TME is too simplistic. Various studies have also identified CAFs with tumor-restraining properties[3,4]. The arrival of single cell genomics has uncovered CAFs as one of the most heterogeneous cell populations within the TME consisting of myofibroblastic or extracellular matrix (ECM) producing CAFs, antigen-presenting CAFs and inflammatory CAFs [5]. Despite increasing knowledge regarding the functions and heterogeneity of CAFs, little is known about the origin of CAFs and whether the different sources of fibroblasts explain the observed heterogeneity. A number of hypotheses have been postulated regarding the cell-of-origin of CAFs, including recruitment of bone marrow (BM) or adipose tissue-derived (AT) mesenchymal stem cells (BM-MSCs or AT-MSCs), epithelial-to mesenchymal transition (EMT) of tumor cells, trans-differentiation of endothelial cells or pericytes and recruitment and activation of local, tissue-resident fibroblasts[2]. Most of these hypotheses have been tested in *in vitro* co-culture settings or in studies involving co-injections of CAF precursors with tumor cells in mice[6–12]. However, assessing the ability of established tumor cells to induce CAF-like behavior in a recipient cell *in vitro* does not mean that these two cell types meet, co-exist or communicate with each other in a similar way during *de novo* tumorigenesis *in vivo.* Only few studies have examined the origin of fibroblasts in more physiological settings. Through a series of *in vivo* transplantations LeBleu and colleagues showed that fibroblasts present in fibrotic kidney disease consisted of locally activated fibroblasts, BM-MSCs and trans-differentiated endothelium-derived fibroblasts[13]. In addition, bone marrow transplantations revealed the existence of a population of bone marrow-derived PDGFRα-negative CAFs in the MMTV-PyMT mouse model of breast cancer[14].

Here, we show through a series of complementary transplantation techniques using genetically engineered mouse models (GEMMs) of invasive lobular breast cancer (ILC) and triple negative breast cancer (TNBC), that the heterogeneous population of CAFs in these tumors originate almost entirely from local, tissue-resident normal fibroblasts (NFs). Assessment of early, progressive and end-stage tumors showed involvement of local, tissue-resident NFs at all stages of tumor development. Single-cell transcriptomics of murine ILC lesions during tumor development revealed the simultaneous disappearance of NFs and appearance of CAFs. Consistent with single-cell transcriptomics datasets of pan-tissue fibroblasts[15] and studies of fibroblasts in normal human breast tissue[16] we identified two subsets of NFs that could be distinguished from each other by their CD26 expression. CD26 (also known as Dpp4) is a T-cell co-stimulatory molecule with peptidase activity and is involved in the deactivation of various hormones and cytokines[17–19]. CD26+ fibroblasts have been linked to scar formation and fibrosis-related diseases[20–23]. Using functional *in vitro* assays and *in vivo* analyses, we showed that CD26+ NFs are predisposed to become inflammatory CAFs (iCAFs), whereas CD26- NFs give rise to myofibroblastic CAFs (myCAFs). CD26+ NFs enhanced the invasive properties of tumor cells via matrix metalloproteinase (MMP) activity and co-cultures of CD26+ NFs and tumor cells recruited CD11b+ myeloid cells in a CXCL12-dependent manner. Taken together, our data show that tissue-resident mammary fibroblasts contribute to the heterogeneous population of CAFs in breast cancer and that CD26+ NFs are at the origin of pro-tumorigenic iCAFs.

## Results

### TNBC and ILC contain CAFs at all stages of tumor development but differ in their relative contents

To investigate and compare the contribution of CAFs to the TME of different breast cancer subtypes we used our clinically relevant GEMMs of BRCA1-deficient TNBC and E-cadherin-deficient ILC. BRCA1-deficient TNBC can be modelled *in vivo* by Cre-conditional, mammary gland-specific loss of BRCA1 and P53 (*WapCre;Brca1^F/F^;Trp53^F/F^*, in short WB1P) alone or in combination with Myc overexpression (*WapCre;Brca1^F/F^;Trp53^F/F^;Col1a1^invCAG-Myc-IRES-Luc^*, in short WB1P-Myc)[24]. ILC was likewise modelled by Cre-mediated mammary-specific loss of E-cadherin and PTEN (*WapCre;Cdh1^F/F^;Pten^F/F^*, in short WEPtn), or by E-cadherin loss with mutant PIK3CA^H1047R^ overexpression (*WapCre;Cdh1^F/F^;Col1a1^invCAG-Pik3caH1047R-IRES-Luc^*, in short WEH1047R)[25]. Histopathological analysis of the resulting mammary tumors showed that TNBC and ILC differed greatly in their CAF content. Sections of end-stage tumors from each of the four breast cancer models and normal mammary glands were stained for fibroblasts using antibodies against PDGFRβ, αSMA and vimentin; for collagen fibers by Masson Trichrome and for epithelial cells using antibodies against E-cadherin and EpCAM (**figure 1A**). ILC lesions contained considerably more fibroblasts than TNBC lesions. Furthermore, the analysis of early, advanced and end-stage tumors isolated from the four mouse models showed a distinct, but remarkably stable presence of fibroblasts at all stages of tumor development (**figure 1B**). The observed differences in CAF content might be a consequence of tumor-specific interactions between CAFs and tumor cells or differences in the cells-of-origin of CAFs. Since little is known about the cells-of-origin of CAFs during *de novo* tumorigenesis *in vivo*, we set out to determine which precursor cells give rise to CAFs in ILC and TNBC.

**Figure 1:**
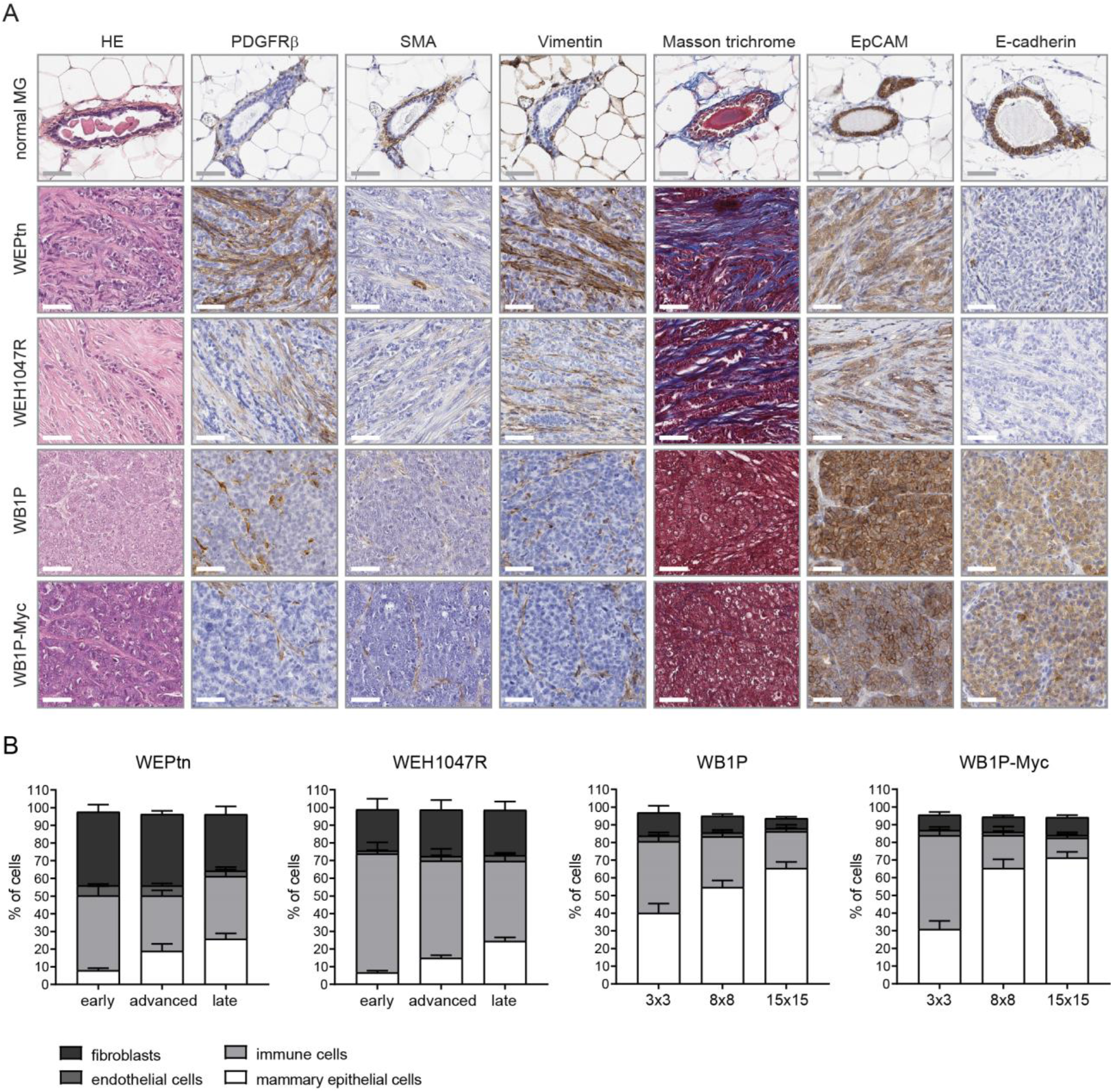
TNBC and ILC differ in histology and quantity of CAFs. A) HE and immunohistochemical stainings of non-tumor bearing control mammary glands and tumors derived from ILC mouse models (WEPtn and WEH1047R) and TNBC mouse models (WB1P and WB1P-Myc) for fibroblast markers (platelet-derived growth factor receptor beta: PDGFRβ, alpha smooth muscle actin: SMA, vimentin), collagen (Masson trichrome) and epithelial cells (epithelial cell adhesion molecule: EpCAM, E-cadherin). Scale bars are 50 um. B) Flow cytometry analysis of tumor composition at indicated time points in WEPtn, WEH1047R, WB1P and WB1P-Myc mice (n=6 per time point). Mammary epithelial cells (tumor cells) were defined as EpCAM+/CD49f+/CD31-/CD45-/PDGFRβ-. Endothelial cells were defined as CD31+/CD49f+/EpCAM-/CD45-. Immune cells were defined as CD45+/CD31-/EpCAM-/CD49f-. Fibroblasts were defined as EpCAM-/CD49f-/CD45-/CD31-/PDGFRβ+.

### Mammary tissue-resident cells contribute to pool of CAFs in breast cancer

We designed a series of complementary transplantation studies to investigate whether CAFs in ILC and TNBC originate from tumor cells that underwent EMT, recruitment of bone marrow-derived MSCs and/or recruitment of local, tissue-resident fibroblasts. (**figure 2A**). Transplantation of pre-neoplastic mouse mammary epithelial cells (MMECs) derived from our ILC and TNBC mouse models into the cleared fat pads of *mTmG* (membrane-tdTomato-membrane-eGFP) recipient mice with ubiquitous expression of cell membrane-localized tdTomato resulted in *de novo* mammary tumor formation.

**Figure 2:**
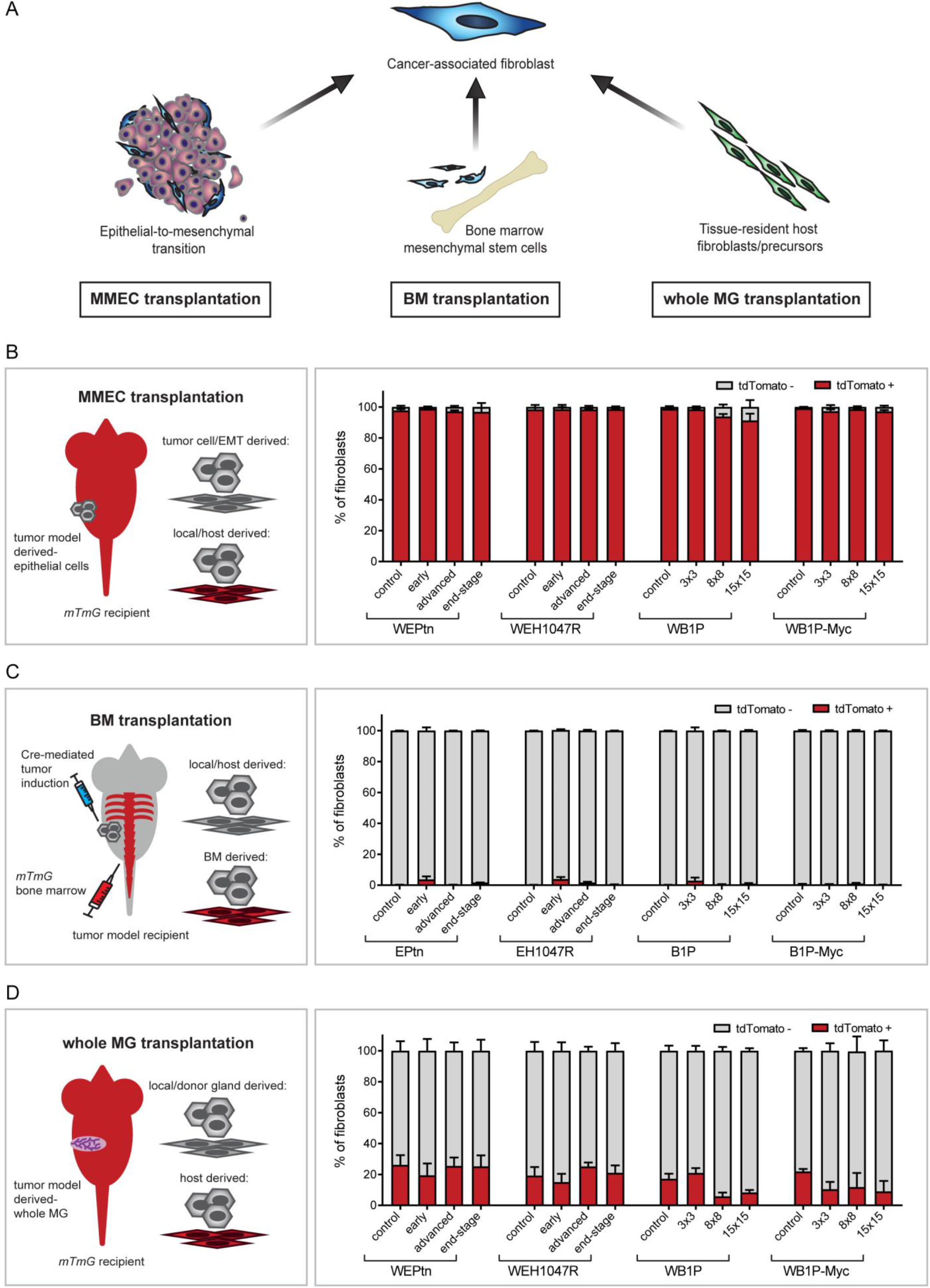
Nearly all CAFs in TNBC and ILC originate from mammary tissue-resident fibroblasts. A) schematic overview of most common CAF-cell-of-origin hypotheses and the transplantation approaches used to validate them. B) Schematic representation of mouse mammary epithelial cell (MMEC) transplantation approach and the results of transplanting pre-neoplastic mammary fragments of WEPtn, WEH1047R, WB1P or WB1P-Myc into the cleared mammary fat pads of tdTomato-positive (mTmG) recipient mice. Tumors were analyzed at early (3×3 mm), advanced (8×8 mm) or end-stage (15×15 mm) by flow cytometry to determine the fraction of tdTomato+ and tdTomato-fibroblasts. Mice transplanted with littermate control, non-neoplastic tissue were used as controls. C) Schematic representation of bone marrow transplantation experiment. tdTomato-positive bone marrow from mTmG mice was transplanted in lethally irradiated mice. Three weeks post transplantation, tumors were induced via intraductal injection of lentivirus expressing Cre-recombinase. Resulting tumors were analyzed at indicated time points by flow cytometry to determine the fraction of tdTomato+ and tdTomato-fibroblasts. Mice that were intraductally injected with PBS and therefore did not develop tumors were used as controls. D) Schematic representation of whole mammary gland transplantation and its results. Pre-neoplastic and littermate control 3^rd^ mammary glands were harvested from WEPtn, WEH1047R, WB1P and WB1P-Myc mice and transplanted in mTmG mice. Tumors and controls were harvested at indicated time points and analyzed by flow cytometry to determine the fraction of tdTomato+ and tdTomato-fibroblasts. All experiments in panels B, C and D were performed with a minimum of 5 mice per time point and in these experiments fibroblasts were defined as EpCAM-/CD49f-/CD45-/CD31-/PDGFRβ+.

Flow cytometry analysis of early, advanced and end-stage tumors showed that nearly all CAFs (EpCAM-/CD45-/CD31-/CD49f-/PDGFRβ+ cells, see **supplemental figure 1** and **2** for gating strategy) expressed tdTomato and were therefore host-derived (**figure 2B**). This does not exclude the possibility that EMT did not occur in these tumors, as all tumors, independent of tumor stage, contained on average 11.08% (± 5.5%) tdTomato-negative cells that lacked expression of epithelial markers (**supplemental figure 3A,B**). Interestingly, these cells did not express the fibroblast-associated marker PDGFRβ (**supplemental figure 3A**). Isolation and culture of EpCAM-positive tumor cells, EpCAM-negative tumor cells and tdTomato-positive CAFs derived from the same WB1P-Myc tumors showed that both EpCAM-positive and EpCAM-negative tumor cells lacked the ability to remodel collagen and gave rise to mammary tumors upon orthotopic re-transplantation. In contrast, tdTomato-positive CAFs were able to remodel collagen *in vitro* and did not lead to tumor formation *in vivo* (**supplemental figure 3C-E**).

Previous reports have shown that bone marrow-derived mesenchymal stem cells (BM-MSCs) can contribute to the pool of CAFs in breast cancer[26,10,14,12]. However, most studies have used xenograft models or co-injections of tumor cells with cultured BM-MSCs. Here, we performed bone marrow (BM) transplantations in our breast cancer mouse models to assess the contribution of BM-MSCs to the pool of CAFs. By using BM derived from *mTmG* donor mice, successful engraftment and BM-derived CAFs could be monitored by their tdTomato expression. Three weeks after BM engraftment of *Cdh1^F/F^;Pten^F/F^* (EPtn), *Cdh1^F/F^;Col1a1^invCAG-^ P^ik3caH1047R-IRES-Luc^* (EH1047R), *Brca1^F/F^;Trp53^F/F^* (B1P) or *Brca1^F/F^;Trp53^F/F^;Col1a1^invCAG-Myc-IRES-Luc/+^* (B1P-Myc) female mice, tumors were induced by intraductal delivery of Cre-encoding lentivirus. Analysis of the resulting mammary tumors showed that nearly all CAFs in the TNBC and ILC models were tdTomato-negative (**figure 2C**). Some tdTomato-positive CAFs were observed in early lesions (EPtn: 3.6% ± 4.1. EH1047R: 4.6% ± 2.1. B1P: 2.8% ± 3.6. B1P-Myc: 0.5% ± 0.6), but these numbers were not statistically significant, suggesting that BM-MSCs were not actively recruited during mammary tumorigenesis. BM harvested from tumor-bearing mice confirmed successful engraftment of donor BM and tdTomato expression in BM-MSCs (**supplemental figure 4**).

Since these findings contradict previous reports based on the PyMT breast cancer mouse model[14], we questioned whether BM-MSC-derived CAFs might have lost tdTomato expression during their recruitment and differentiation and repeated this experiment using *Cdh1^F/F^;Pten^F/F^;mTmG* mice transplanted with BM from wild-type FVB/n mice (**supplemental figure 5A**). Now, the transplanted BM lacked fluorescence and the recipient mouse expressed the *mTmG* Cre-conditional reporter that replaced tdTomato expression with GFP in tumor cells (**supplemental figure 5B,C**). Again, BM-MSCs were not the main contributors to the population of CAFs, as the majority of the CAFs in the resulting mammary tumors were tdTomato-positive (**supplemental figure 5D,E**).

To determine whether breast CAFs originate from tissue-resident fibroblasts or their precursors, we performed whole mammary gland transplantations[27]. Transplanting whole pre-neoplastic mammary glands including the nipple and associated skin isolated from WEPtn, WEH1047R, WB1P and WB1P-Myc mice into *mTmG* recipients allowed for discrimination between gland- and host-derived cells. Analysis of tumors arising in the transplanted mammary glands revealed that the majority of CAFs were tdTomato-negative and therefore descendants of the transplanted gland rather than the host (**figure 2D**). Although a substantial number of CAFs were tdTomato-positive, this was also observed in recipients transplanted with control mammary glands from WapCre-negative littermates, indicating a surgery-induced effect (**figure 2D**). Nevertheless, the organization of the transplanted glands resembled endogenous gland architecture and cellular distribution (**supplemental figure 6**). In summary, these transplantation-based *in vivo* studies showed that CAFs in TNBC and ILC largely originate from tissue-resident fibroblasts or precursors.

### Single-cell transcriptomics reveal NF and CAF dynamics during tumor development

Recent advances in single-cell transcriptomics have highlighted CAF heterogeneity and the existence of distinct CAF subpopulations in various cancer types[28,5,29]. Having shown that CAFs originate from mammary tissue-resident cells, we decided to investigate the heterogeneity of normal mammary fibroblasts (NFs) and their transition into CAFs during *de novo* tumorigenesis. For this purpose, we focused on ILC as these tumors show a large infiltrate of CAFs and little is known about their role in ILC development and progression. ILC mammary lesions were induced by intraductal injection of Cre-encoding lentivirus in EPtn mice and CAFs were isolated from these tissues 6-, 12- or 18-weeks post-injection. Together with control NFs, these CAFs were subjected to single-cell transcriptomics, which revealed a progressive disappearance of NF clusters and a concomitant appearance of CAF clusters during tumor progression (**figure 3A,B**). Gene ontology analysis of the top-25 differentially expressed genes that define the clusters identified specific functions for each CAF cluster such as immune-modulation (cluster 2, iCAFs) and ECM-production (cluster 4, myCAFs) (**figure 3C-E**). Using these top-25 differentially expressed genes we generated an iCAF and myCAF gene expression signature (**supplemental figure 7A**) to investigate in other datasets.

**Figure 3:**
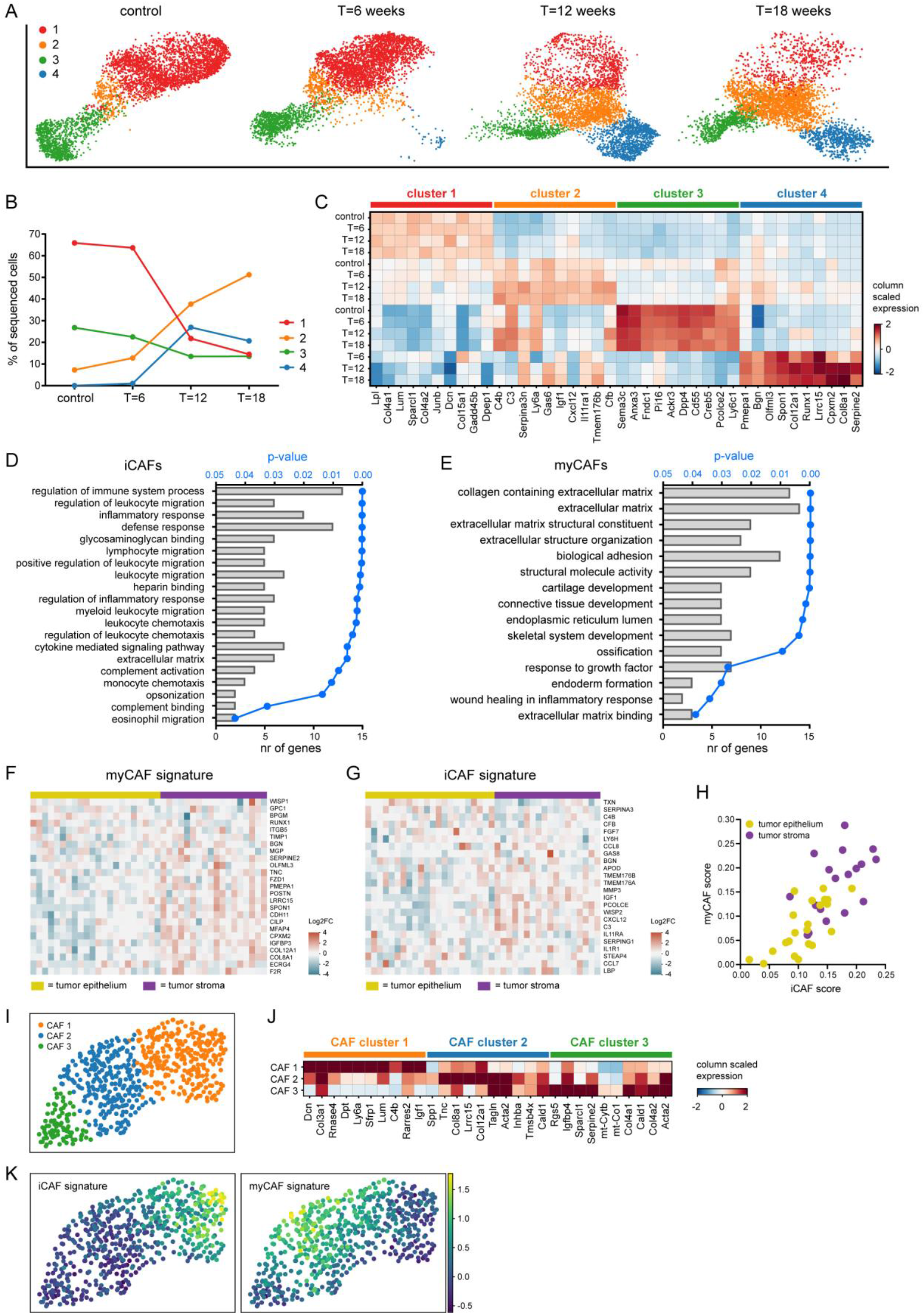
Single-cell transcriptomics reveal dynamics in NFs and CAFs during tumor development. A) Fibroblasts were isolated from control, non-tumor bearing mice and ILC-tumor bearing mice (*Cdh1^F/F^;Pten^F/F^* mice) 6, 12 or 18 weeks after intraductal lenti-Cre injection to initiate tumor formation (n=2 mice per time point). Fibroblasts were defined as EpCAM-/CD45-/CD31-/CD49f-cells, sorted by FACS and subjected to single-cell RNA sequencing using the 10X Genomics platform. The data was analyzed using scanpy and represented as UMAP plots for each time point. B) Dynamics of the 4 fibroblast clusters during tumor development. Number of fibroblasts in each cluster is expressed as percentage of all sequenced cells of the indicated time points. C) Matrix plot showing the top-10 genes that define each cluster and their expression at each time point. D) Gene ontology analysis of top-25 genes that define CAF cluster 2 (immune-modulating CAFs, iCAFs). E) Gene ontology analysis of top-25 genes that define CAF cluster 4 (myofibroblastic CAFs, myCAFs). F) iCAF gene expression signature in laser-microdissected human ILC samples (n=17) separated into tumor epithelium (marked in yellow) and tumor stroma (marked in purple). G) myCAF gene expression signature in human ILC. H) Single sample gene set enrichment analysis of iCAF and myCAFs scores in human ILC tumor epithelium and tumor stroma. I) Reclustering of fibroblasts (cluster 5 from supplemental figure 7E) from single-cell transcriptomics analysis of end-stage WB1P and WB1P-Myc tumors. J) Matrix plot showing the top-10 genes that define each CAF subcluster. K) UMAP plot of the ILC iCAF and myCAF signatures within the single-cell transcriptomics dataset of CAFs from WB1P and WB1P-Myc.

Recently, Gómez-Cuadrado *et al.* investigated the characteristics of the non-immune stroma in human ILC and IDC by laser-capture microdissection and transcriptomic analysis[30]. We interrogated this dataset to determine whether murine iCAF and myCAF gene signatures were also expressed in human ILC and IDC. Various iCAF- and myCAF-related genes were expressed in human ILC (**figure 3F, G**) and IDC stroma (**supplemental figure 7B,C**). Furthermore, single sample gene set enrichment analysis showed that the iCAF and myCAF signature scores were enhanced in tumor stroma compared to tumor epithelium, indicating that similar iCAF and myCAF populations exist in human ILC (**figure 3H**) and IDC (**supplemental figure 7D**).

To determine if iCAFs and myCAFs are also present in TNBC, we performed single-cell transcriptomics on end-stage mammary tumors from WB1P and WB1P-Myc mice. Based on marker gene expression we identified various clusters of tumor cells, immune cells, endothelial cells and CAFs (**supplemental figure 7E**). The expression of *Col1a1* and *Col1a2* was used to identify the cluster of CAFs in WB1P and WB1P-Myc tumors (**supplemental figure 7F**). Reclustering of the CAFs yielded three subclusters with distinct differences in gene expression (**figure 3I,J**). Subcluster 1 expressed *Ly6a, C4b* and *Igf1*, which are also expressed by ILC iCAFs, whereas subclusters 2 and 3 expressed *Lrrc15, Col8a1* and *Serpine2*, which are also upregulated in ILC myCAFs. As expected, analysis of the ILC iCAF and myCAF signatures within TNBC-derived CAFs showed that both WB1P and WB1P-Myc mammary tumors contain CAFs with iCAF and myCAF signatures (**figure 3K**).

Similar CAF subpopulations have been described in human pancreatic ductal adenocarcinoma (PDAC) and PDAC mouse models [5]. Interestingly, these PDAC iCAF and myCAF signatures showed substantial overlap with the iCAF and myCAF clusters in ILC (**supplemental figure 8A-D**). We did not find a distinct ILC CAF cluster that resembled the antigen-presenting CAFs (apCAFs) observed in PDAC, although some cells in the ILC iCAF cluster appeared positive for the PDAC apCAF signature (**supplemental figure 8A**). Comparison of gene expression profiles from PDAC iCAFs and myCAFs with ILC iCAFs and myCAFs beyond the signature showed significant correlations in both up- and downregulated genes in iCAFs and myCAFs from ILC and PDAC (**supplemental figure 8E, F**), indicating that, independent of cancer type, these CAF subtypes emerge during tumor development.

Mammary NFs could be separated into CD26- (cluster 1) and CD26+ (cluster 3) cells (**figure 4A**). The presence of two distinct fibroblast populations that could be distinguished from each other by CD26 expression has previously been reported in normal human and mouse mammary tissue[16,31]. Gene ontology analysis of the differentially expressed genes that define CD26- and CD26+ NFs primarily revealed functions related to ECM production and maintenance (**supplemental figure 9A-B**). Further investigation of the genes annotated to the common GO term ‘Extracellular matrix’ revealed differences between CD26- and CD26+ NFs (**supplemental figure 9C**). CD26- NFs primarily express collagens (*Col15a1, Col18a1, Col4a1, Col4a2* and *Col5a3)* and proteins involved in collagen fibril formation, which have been previously linked to inhibition of angiogenesis, metastasis and tumor suppression (*Dcn*, *Lum*, *Sparcl1* and *Anxa5*)[32–34]. In contrast, CD26+ NFs predominantly express fibronectin (*Fn1*), fibrillin (*Fbn1)* and elastic fiber-related ECM molecules (*Emilin2)* that have been associated with enhanced angiogenesis and metastasis (*Cd248*, *Emilin2*, *Dpt*)[35–38].

**Figure 4:**
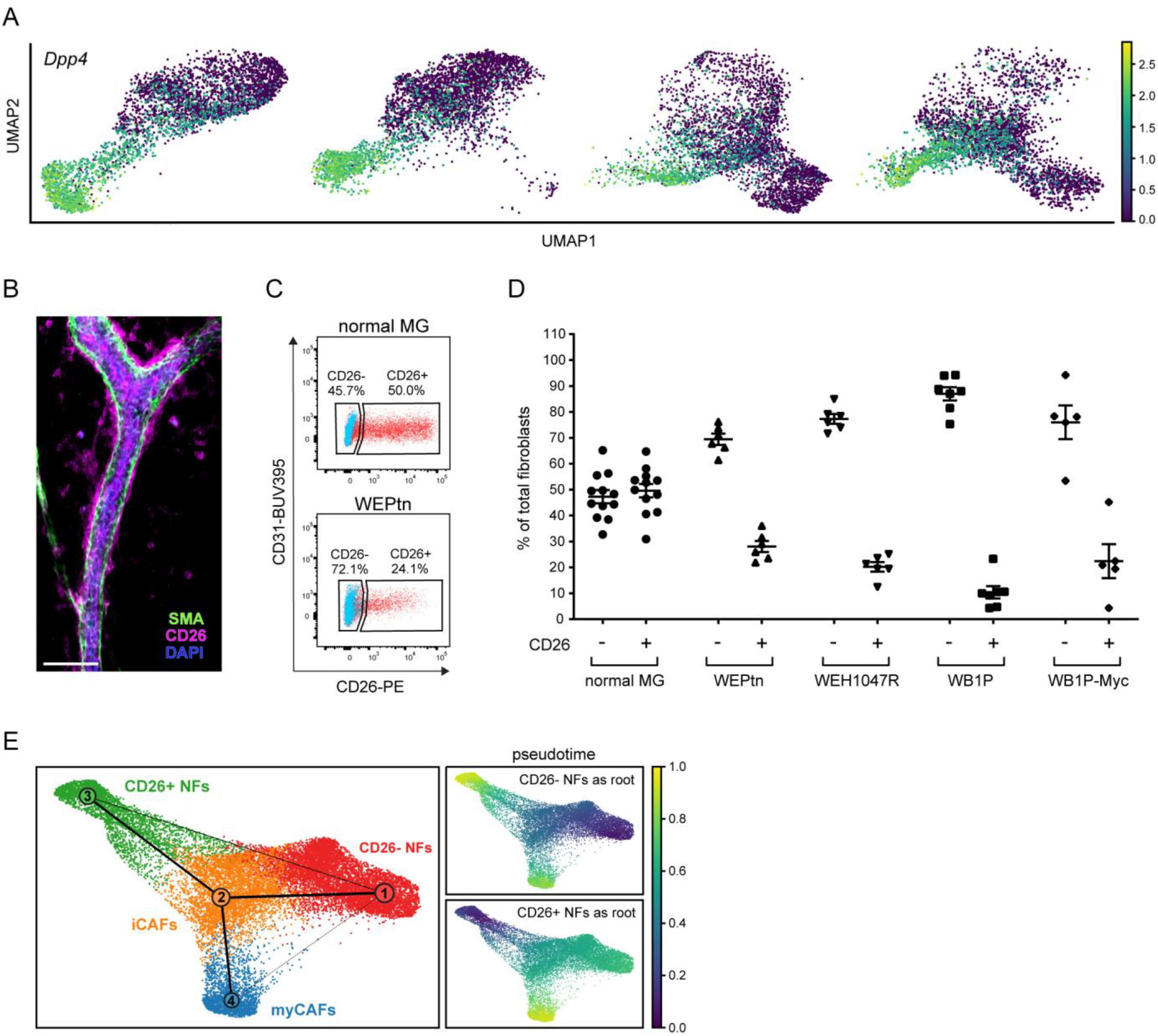
Ratio of CD26- and CD26+ fibroblasts change during tumor development and both CD26- and CD26+ NFs contribute to CAF populations. A) UMAP plots of single-cell transcriptomics data showing *CD26 (Dpp4)* expression. B) Whole mount analysis of normal mammary gland stained for CD26 (in magenta) and smooth muscle actin (SMA, in green) and dapi (in blue). Scale bar is 50 um. C) Flow cytometry analysis of normal mammary gland and WEPtn-derived ILC gated on fibroblasts (EpCAM-/CD45/-CD31-/CD49f-cells) shows the presence of CD26- and CD26+ fibroblasts. Shown in blue is fluorescence-minus-one (FMO) control and shown in red is full stained sample. FMO control was used to set gates. D) Percentage of CD26- and CD26+ fibroblasts in control mammary glands (n=12 mice) and end-stage tumors derived from WEPtn (n=6 mice), WEH1047R (n=6 mice), WB1P (n=7 mice) and WB1P-Myc mice (n=5 mice). E) Partition-based graph abstraction (PAGA) trajectory analysis of single-cell transcriptomics data indicates trajectory of NF clusters towards CAF clusters. Pseudotime was plotted using CD26- NFs as a starting point (root) and CD26+ NFs as a starting point.

The presence of CD26- and CD26+ NFs in the mammary gland was confirmed by whole mount analysis and flow cytometry (**figure 4B,C**). Interestingly, a decrease in CD26+ fibroblasts was observed within the stroma of mammary tumors compared to wild type mammary glands, confirming the absence of CD26 in myCAFs and low CD26 expression in iCAFs within the single-cell transcriptomics dataset (**figure 4A,D**). This shift in the ratio of CD26- and CD26+ fibroblasts was observed in both ILCs and TNBCs and became more apparent as tumors progressed (**figure 4D and supplemental figure 10**). To determine if CD26- and CD26+ NFs contribute to specific CAF populations, we performed a trajectory analysis, which showed that both CD26- NFs and CD26+ NFs can transition into iCAFs and subsequently into myCAFs. In addition, a small fraction of CD26- NFs seemed to directly transition into myCAFs (**figure 4E**). These data suggest that both CD26- and CD26+ NFs contribute to iCAF and myCAF populations in mammary tumors. Whether both CD26+ and CD26- NF-derived iCAFs can form myCAFs remains unclear from this analysis.

### CD26+ NFs enhance tumor cell invasion

Next, we set out to investigate whether functional differences exist between CD26- and CD26+ NFs. Primary mammary CD26- and CD26+ NFs were harvested and cultured to determine whether they differentially affected tumor cell behavior. Both NF populations displayed spindle-shaped cell morphology and parallel alignment associated with cultured fibroblasts (**supplemental figure 11A**). In a transwell setting, CD26+ NFs displayed preferential recruitment towards ILC derived tumor cells, whereas TNBC cells recruited both CD26- and CD26+ NFs (**figure 5A-C**). The preferential recruitment of CD26+ NFs towards ILC-derived tumor cells was not dependent on CD26 enzymatic activity, as CD26 inhibitors did not affect recruitment (**supplemental figure 11B**). Interestingly, when CD26- and CD26+ NFs were mixed prior to plating, both NF populations migrated towards ILC tumor cells, suggesting crosstalk between CD26- and CD26+ NFs in this experimental setting (**supplemental figure 11C, D**). CAFs have previously been shown to enhance the metastatic potential of tumor cells[2,39–44]. To determine whether CD26- or CD26+ NFs specifically promoted tumor cell migration, they were subjected to an organotypic invasion assay to measure migration of tumor cells into a matrix containing collagen and basement membrane extract (BME) with or without NFs. Although both CD26- and CD26+ NFs promoted tumor cell invasion, migration into the matrix was more strongly enhanced by CD26+ NFs compared to CD26- NFs. This effect was most prominent when CD26+ NFs were combined with ILC-derived tumor cells (**figure 5D, E**), but was also observed with TNBC-derived tumor cells (**supplemental figure 11E, F**). Together, these findings indicate that CD26+ NFs are at the origin of pro-tumorigenic CAFs.

**Figure 5:**
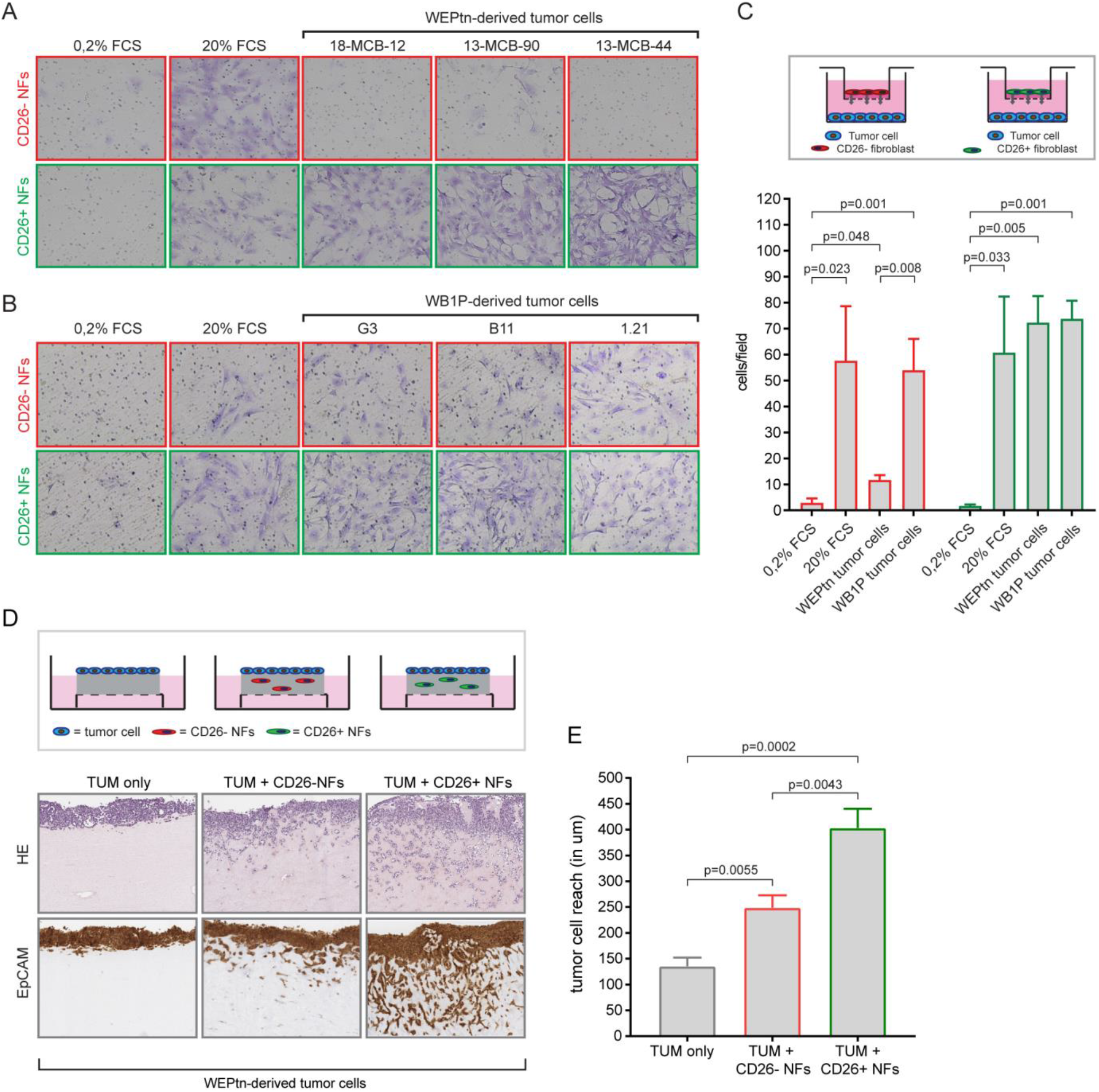
CD26+ NFs are recruited towards tumor cells and induce invasiveness. A) Representative result of fibroblast migration towards tumor cells. Transwell assays were used to assess the recruitment of CD26- and CD26+ fibroblasts towards ILC-derived tumor cells (panel A) and TNBC-derived tumor cells (panel B). Tumor cells were plated in the bottom compartment and fibroblasts were plated in matrigel-coated inserts. 24 hours after plating the bottom-side of the inserts were fixed and stained with crystal violet to visualize the migrated cells. All assays were done in low serum conditions (0.2% FCS). 20% FCS and 0.2% FCS alone were used as positive and negative controls, respectively. C) Quantification of transwell assays (n=7 independent experiments with similar outcome). Statistical significance was determine using student’s t-test. D) Representative result of organotypic invasion assay, in which collagen-containing gels are loaded with CD26- or CD26+ NFs or left empty. Tumor cells are plated on top of these gels and invasion into the gels is assessed after one week. Gels are processed as HE slides or stained for EpCAM to visualized tumor cells. E) Quantification of organotypic invasion assays based on EpCAM staining (n=5 independent experiments with similar outcome). Invasion was measured in um from top of the gel to the invasive front of the tumor cells. Statistical significance was determine using student’s t-test.

### CD26- and CD26+ NFs are predisposed to become myCAFs and iCAFs, respectively

Next, we determined if both CD26- and CD26+ NFs were able to adopt an iCAF-like gene signature, as suggested by the trajectory analysis. For this purpose, we cultured primary CD26- and CD26+ NFs in conditioned medium (CM) derived from WEPtn-derived tumor cells and compared their gene expression profiles with those of NFs cultured in control medium (**figure 6A**). Hierarchical clustering of the top-100 differentially expressed genes showed that CD26- and CD26+ NFs responded differently to the same tumor CM (**supplemental figure 12A**).

**Figure 6:**
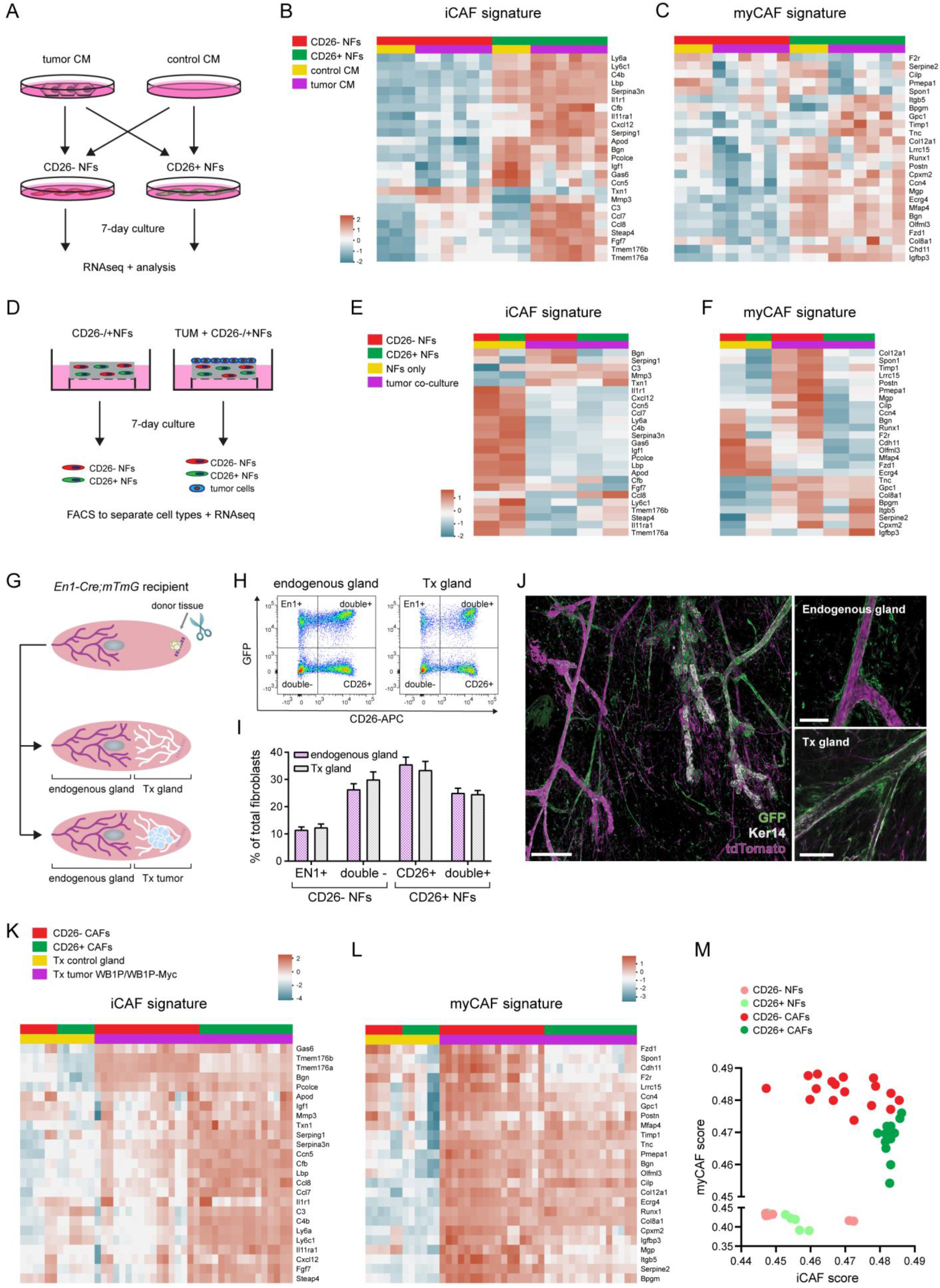
CD26- NFs are predisposed to become myCAFs, whereas CD26+ NFs become iCAFs. A) Schematic overview of experimental set-up in which CD26- NFs and CD26+ NFs are cultured for 7 days in conditioned medium (CM) derived from 6 independent WEPtn-derived ILC tumor cell lines followed by RNA sequencing. B) Transcriptomics analysis of the iCAF gene signature in CD26- NFs and CD26+ NFs cultured in control CM or tumor CM. C) Analysis of the myCAF gene signature in CD26- NFs and CD26+ NFs cultured in control CM or tumor CM. D) Schematic overview of experimental set-up in which CD26- and CD26+ NFs were cultured in collagen-containing matrix alone or in the presence of tumor cells (2 WEPtn-derived tumor cell lines) for 7 days. After 7 days the cell types were separated by FACS and sequenced. E and F) Analysis of the iCAF (panel E) and myCAF (panel F) gene signatures in CD26- and CD26+ NFs cultured alone or in presence of tumor cells. G) Schematic representation of MMEC transplantations in *En1-Cre;mTmG* in which control littermate or pre-neoplastic donor tissue is placed at the dorsal tip of the recipient mammary gland of three-week old *En1-Cre;mTmG* mice. H) Representative flow cytometry plots of the distribution of mammary fibroblasts in *En1-Cre;mTmG* control mice and *En1-Cre;mTmG* mice transplanted with control mammary tissue. Fibroblasts were defined as EpCAM-/CD49f-/CD31-/CD45-cells. I) Quantification of indicated fibroblasts populations in endogenous mammary glands of *En1-Cre;mTmG* mice and mammary glands with transplanted control donor tissue (n=6 mice per group). Statistical significance was determined using student’s t-test. No statistical differences found between control and transplanted (Tx) glands for the different populations. J) Whole mount analysis of transplanted gland showing the endogenous mammary ductal structure in magenta (tdTomato) and the outgrown transplanted donor tissue in white (stained for keratin 14). Fibroblasts from the Engrailed1 lineage are shown by their GFP expression in green. Zoom images on the right show GFP+ fibroblasts surrounding both endogenous and transplanted mammary ducts. Scale bar of overview image is 200 um. Scale bar in image of endogenous gland is 50 um and scale bare in image of Tx gland is 100 um. K and L) Transcriptomics analysis and iCAF (panel K) and myCAF (panel L) gene signatures of CD26- and CD26+ NFs and CAFs isolated from *En1-Cre;mTmG* mice transplanted with control or pre-neoplastic mammary tissue from WB1P and WB1P-Myc mice. Fibroblasts were isolated by FACS from control transplanted mice (n=3) and end-stage tumors (WB1P n=5, WB1P-Myc n=4). M) single sample gene set enrichment analysis (Singscore) of the iCAF and myCAF signatures in CD26- and CD26+ NFs and CAFs isolated from *En1-Cre;mTmG* mice.

KEGG pathway analysis showed enrichment in pathways associated with cytokine-cytokine receptor interaction and chemokine signaling in CD26+ NFs cultured in CM compared to control CD26+ NFs. CD26- NFs exposed to tumor CM showed enrichment in pathways associated with tight junction regulation and cellular contractility (**supplemental figure 12B, C**). The KEGG pathway analysis results founded our hypothesis that CD26+ NFs contribute to the population of iCAFs and could indicate that CD26- NFs contribute to the population of myCAFs. To investigate this in more detail, we investigated whether the genes of the iCAF and myCAF signatures from our single-cell transcriptomics dataset were upregulated or enriched in CM-exposed CD26+ and CD26- NFs. As expected, the iCAF gene signature was enriched in the CD26+ NFs cultured in tumor CM (**figure 6B**), further supporting our hypothesis. However, the myCAF signature was not enriched in CD26- or CD26+ NFs cultured in tumor CM (**figure 6C**). Since this experiment was performed in the absence of ECM and direct tumor cell-fibroblast contact, we repeated the experiment by co-culturing tumor cells and NFs in a collagen-rich matrix to better recapitulate the *in vivo* situation (**figure 6D**). Analysis of the CAF signatures within this experimental set-up showed an absence of the iCAF signature in both CD26- and CD26+ NFs co-cultured with tumor cells (**figure 6E**). However, the myCAF signature was enriched in the CD26- NFs co-cultured with tumor cells compared to fibroblasts cultured in the absence of tumor cells. Moreover, the myCAF signature was not apparent in the CD26+ NFs of the same co-culture (**figure 6F**). Taken together, these *in vitro* studies suggest that CD26- NFs are predisposed to become myCAFs whereas CD26+ NFs are predisposed to become iCAFs. Nevertheless, induction of these phenotypes is highly dependent on the culture conditions used. Therefore, we investigated whether this predisposition also exists *in vivo*. Previous reports showed that Engrailed1 (En1) marks a pro-fibrotic lineage of fibroblasts in the skin and that En1-positive skin fibroblasts also express CD26[23]. Therefore, *En1-Cre;mTmG* mice would allow for *in vivo* tracing of GFP-positive CD26+ NFs and tdTomato-positive CD26- NFs during tumor development independent of changes in gene expression. For this purpose we performed MMEC transplantations in *En1-Cre;mTmG* mice (**figure 6G**). However, contrary to the expected results, in the mammary gland not all En1+ NFs expressed CD26 and a substantial amount of CD26+ NFs do not originate from the En1+ lineage (**figure 6H**). MMEC transplantations were performed as described previously, with the exception of clearing of the recipient mammary fat pad (**figure 6G**). By keeping the recipient gland intact and placing donor tissue at the dorsal side of the fourth mammary gland, we ensured minimal perturbations to the recipient gland and the fibroblasts present in that gland. Transplantation of control donor tissue from WapCre-negative *Cdh1^F/F^;Pten^F/F^* mice in *En1-Cre;mTmG* mice verified proper outgrowth and comparable distributions of fibroblasts in transplanted and control glands (**figure 6H-J**). Analysis of the mammary tumors that arose from transplantations of pre-neoplastic mammary donor tissue from WEPtn, WB1P and WB1P-Myc mice in *En1-Cre;mTmG* mice revealed an increase in En1+/CD26- (En1+) and EN1-/CD26- (double-) fibroblasts, a decrease in EN1-/CD26+ (CD26+) fibroblasts and similar levels of EN1+/CD26+ (double+) fibroblasts in tumors compared to controls (**supplemental figure 13A-C**). Transcriptomics analysis and hierarchical clustering of En1+, CD26+, double+ and double-fibroblasts isolated from tumors and control mammary glands showed that clustering did not depend on En1 status, but rather on CD26 expression, as double- and En1+ fibroblasts clustered together and CD26+ and double+ fibroblasts clustered together (**supplemental figure 13D**). For simplicity we collectively refer to En1+ and double-fibroblasts as CD26-fibroblasts and CD26+ and double+ fibroblasts as CD26+ fibroblasts. Analysis of the myCAF and iCAF signatures within this dataset showed that the iCAF signature was enhanced in CD26+ CAFs, whereas the myCAF signature was predominantly expressed by CD26-CAFs (**figure 6K, L**). Single sample gene set enrichment analysis verified that CD26+ CAFs express genes associated with the iCAF signature whereas CD26-CAFs express myCAF-associated genes (**figure 6M**). Nevertheless, several genes within the myCAF signature were also expressed by CD26+ CAFs, indicating that these CAFs may also have some myofibroblastic properties. Taken together, these results show that an *in vivo* predisposition exists for CD26- NFs to become myCAFs, whereas CD26+ NFs become iCAFs and that this predisposition is not dependent on En1 status.

### CD26+ NFs co-cultured with tumor cells secrete CXCL12 to recruit monocytes

Our single-cell transcriptomics showed that iCAFs express various cytokines (*Ccl2, Ccl7, Ccl8, Cxcl2* and *Cxcl12)* that play a role in the recruitment of myeloid cells (**figure 7A,B**). Cytokine array analysis of CM derived from CD26- and CD26+ NFs co-cultured with tumor cells revealed that the co-culture of CD26+ NFs and tumor cells released more CXCL2 and CXCL12 than the co-culture of CD26- NFs and tumor cells. Conversely, co-cultures of CD26- NFs and tumor cells secreted more TNFα and CD54 compared to co-cultures of CD26+ NFs and tumor cells. (**figure 7C, supplemental figure 14A,B**). Secretion of CXCL12 from fibroblasts has previously been linked with enhanced tumor growth, angiogenesis and recruitment of T-regulatory cells [29,45]. To determine if CD26+ NFs and their release of cytokines upon co-culture with tumor cells are involved in the recruitment of immune cells, we harvested splenocytes and monitored their recruitment in transwell assays towards CM derived from NF mono- or co-cultures with tumor cells. We found that CM of CD26+ NFs co-cultured with tumor cells recruited more CD11b+ monocytes than all other mono- and co-cultures, indicating an immune-modulatory function of CD26+ NFs (**figure 7D**). No differences were observed in the ability of the NF mono- or co-cultures to recruit CD3+ T-cells or B220+ B-cells (**supplemental figure 14C, D**). Neutralizing antibodies against CXCL2 and CXCL12 revealed that only inhibition of CXCL12 completely abrogated the recruitment of monocytes (**figure 7D**). Since CXCL12 has also been associated with fibroblast-induced tumor cell migration [42,40,46], we assessed whether CD26+ NF-derived CXCL12 was also responsible for the observed invasion of tumor cells in the organotypic invasion assays (**figure 5D**).

**Figure 7:**
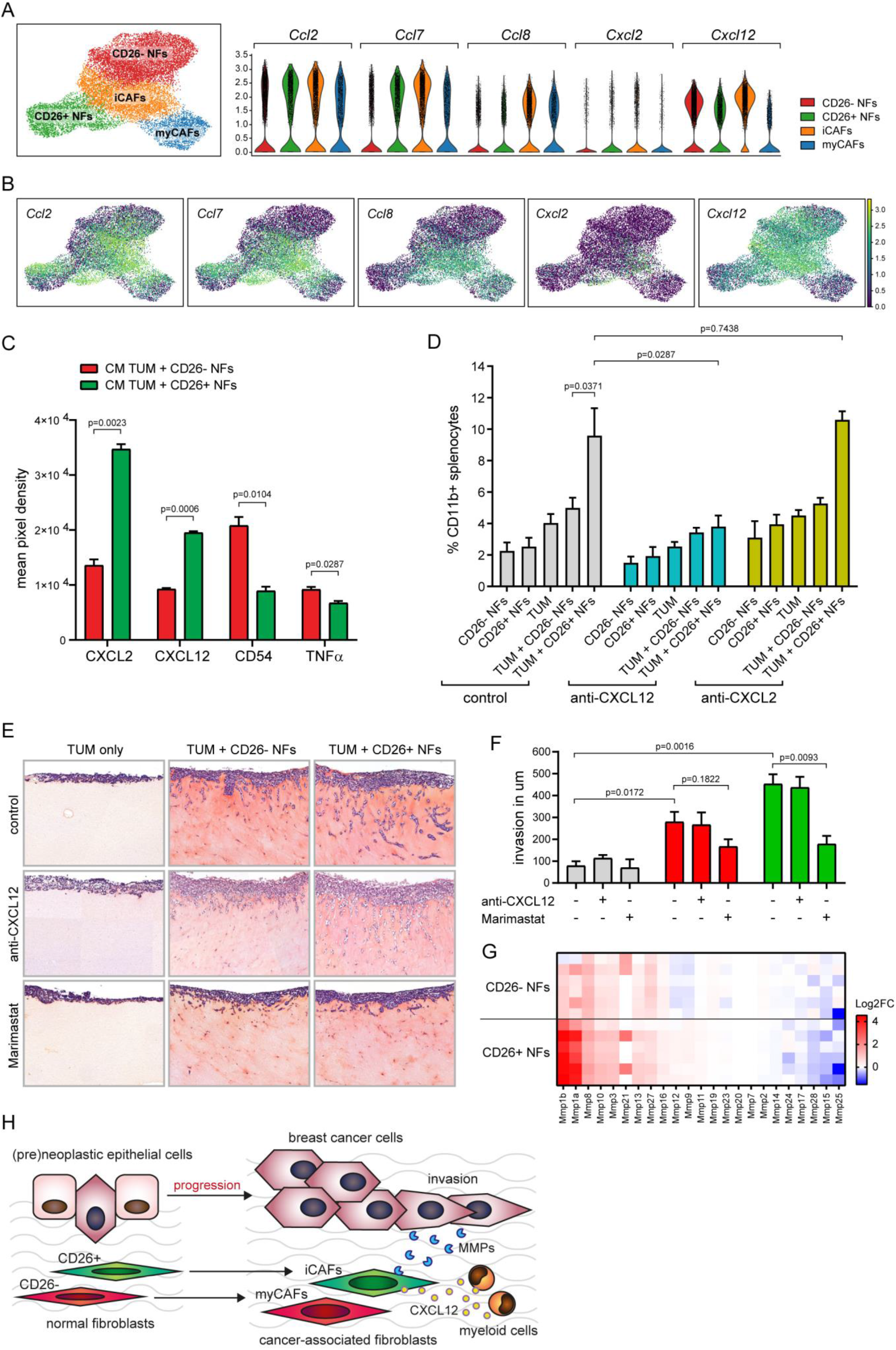
CD26+ NFs recruit CD11b+ myeloid cells in a CXCL12 dependent manner and induce tumor cell invasion via MMPs. A) UMAP plot of single-cell transcriptomics dataset (all time points combined) showing the NF and CAF clusters and the violin plots of the expression levels of *Ccl2, Ccl7, Ccl8, Cxcl2* and *Cxcl12* in each of the clusters. B) UMAP plot representation of data in panel A. C) Significantly up- and downregulated cytokines present in the conditioned medium of indicated co-cultures of tumor cells and fibroblasts as determined by cytokine array. Statistical significance was determine using student’s t-test. D) Results of transwell assays used to investigate recruitment of splenocytes towards the CM of fibroblast and tumor cell mono- or co-cultures in the presence or absence of CXCL2 or CXCL12 neutralizing antibodies. Migrated splenocytes were harvested from the bottom compartment after 24 hours, stained for CD11b, quantified by flow cytometry and displayed as percentage of CD11b+ splenocytes from total live single cells. Results shown are from 3 independent experiments with similar outcome. E) Representative images of organotypic invasion assays in the presence or absence of CXCL12 neutralizing antibodies or the MMP inhibitor Marimastat. F) Quantification of organotypic invasions assays in panel E. Invasion was measured in um from top of the gel to the invasive front of the tumor cells. Statistical significance was determine using student’s t-test. Results are shown from 3 independent experiments with similar outcome. G) Log2 fold change (Log2FC) in expression of MMPs in CD26- and CD26+ NFs cultured in tumor CM compared CD26- and CD26+ NFs cultured in control CM. H) Schematic representation of the transition of CD26- and CD26+ NFs towards myCAFs and iCAFs, respectively during mammary tumorigenesis. CD26+ NFs transform into pro-tumorigenic iCAFs that release CXCL12 to recruit CD11b+ myeloid cells and induce tumor cell migration via MMPs.

Addition of CXCL12 neutralizing antibodies did not affect CD26+ NF-induced tumor cell invasion (**figure 7E, F**). In addition, matrix metalloproteinases (MMPs) have been shown to play an important role in tumor cell migration and invasion[47]. Inhibition of MMPs using marimastat, a broad spectrum MMP inhibitor, blocked tumor cell invasion induced by CD26- and CD26+ NFs (**figure 7E, F**). In line with these results, we found significant changes in gene expression of MMPs in CD26- and CD26+ NFs cultured in tumor CM compared to NFs cultured in control medium. When comparing differential MMP expression in CD26- NFs cultured in tumor CM with CD26+ NFs cultured in tumor CM, we found significant changes in the expression of MMP1, 3, 9, 11, 12, 13 and 28. Two of these MMPs, MMP1 and 9, are drug targets of marimastat, and are significantly increased in CD26+ NFs compared to CD26- NFs (**figure 7G**). Taken together, our results showed that nearly all CAFs in breast cancer originate from tissue-resident mammary fibroblasts and that CD26- NFs are predisposed to become myCAFs, whereas CD26+ NFs are predisposed to become pro-tumorigenic iCAFs, capable of recruiting myeloid cells via CXCL12 secretion and enhancing tumor cell invasion via MMP signaling (**figure 7H**).

## Discussion

Here, we have shown through a series of complementary transplantation techniques using multiple GEMMs, that nearly all CAFs in breast cancer originate from tissue-resident fibroblasts, with little to no contribution from EMT or bone marrow precursors. The lack of bone marrow involvement was surprising, since Raz *et al.* showed *in vivo* recruitment of BM-MSCs in tumors isolated from MMTV-PyMT mice[14]. The differences between our results and those of Raz *et al.* may be due to experimental design. The rapid development of mammary tumors in the PyMT model may create an overlap between the time of BM transplantation and tumor development which could influence recruitment, especially since irradiation can lead to a tissue damage response which BM-MSCs are likely to home to[48]. In our experimental set-up BM engraftment preceded tumor induction by three weeks, reducing the change of interference between the two processes.

Single-cell transcriptomics of fibroblasts at several stages of tumor development have revealed a gradual shift of two normal fibroblast subpopulations into two CAF populations with distinct functions. Throughout this study we used CD26 as a marker to separate the two normal fibroblast populations identified by single-cell transcriptomics. This is consistent with previous reports of CD26+ and CD26-fibroblasts in human and murine mammary tissue [16,31], although other markers may also be used to separate these two fibroblast subtypes. A recent study by Buechler *et al.* generated a single-cell transcriptomics atlas of murine fibroblasts isolated from various tissues. They identified two main populations (*Pi16*+ and *Col15a1+)* present in all investigated tissues[15]. Integration of this data with single-cell transcriptomics data of mammary fibroblasts by Yoshitake *et al.* showed that the *Pi16*+ fibroblasts overlapped with the mammary CD26+ fibroblasts described by Yoshitake *et al*.[31]. In addition, the CD26- NFs overlapped with *Col15a1+* fibroblasts described by Buechler *et al.* Interestingly, Yoshitake *et al.* have found that mammary CD26- and CD26+ fibroblasts displayed population-specific responses to estrogen treatment, indicating different roles of these fibroblasts in mammary gland development and maintenance. These results support our observations of the differential responses of CD26- and CD26+ NFs to tumor CM and co-cultures with tumor cells. CD26 is a known co-stimulatory molecule involved in T-cell activation[17]. In addition, the extracellular domain of CD26 has enzymatic activity that plays an important role in the inactivation of various signaling molecules including incretins, chemokines and cytokines[49]. Our *in vitro* experiments using CD26 inhibitors suggest that CD26 enzymatic activity is not required for the recruitment of fibroblasts towards tumor cells, but further studies are needed to determine the role of CD26 in the transition to iCAFs and their pro-tumorigenic functions.

iCAFs and myCAFs constitute the majority of CAFs found in both ILC and TNBC. Interestingly, these CAF populations showed significant overlap in gene expression with PDAC-derived iCAFs and myCAFs. Others have identified iCAFs and myCAFs in a various malignancies including pancreatic cancer, colorectal cancer, breast cancer and other solid tumors[50,51,5,29,52], suggesting that the emergence of these CAF subtypes is a pan-cancer effect. Surprisingly, the origin of this heterogeneity is understudied. In pancreatic cancer it has been shown that TGF-β and IL1 are tumor-secreted factors that aid in the transformation of pancreatic stellate cells towards a myCAF or iCAF phenotype, respectively[51]. In addition, Miyazaki *et al.* has shown that differential exposure to Wnt drives the transformation of adipose-derived mesenchymal stem cells into iCAFs or myCAFs *in vitro*[7]. However, these studies did not consider normal fibroblast heterogeneity, making it unclear if the observed CAF heterogeneity is a direct consequence of pre-existing heterogeneity within the cells of origin *in vitro* and *in vivo.* We are the first to report a predisposition of two normal mammary fibroblast subtypes to develop into two functionally distinct CAF populations. Using various co-cultures systems we uncovered that CD26+ NFs adopt an iCAF phenotype whereas CD26- NFs are more likely to adopt a myCAF phenotype. However, the observed CAF subtype was highly dependent on the culture system used, emphasizing that caution is needed when drawing conclusions based on *in vitro* assays using highly plastic cells such as fibroblasts. *In vivo* tracing of CD26- and CD26+ fibroblasts in *En1-Cre;mTmG* mice confirmed that CD26-fibroblasts adopt a myCAF signature and CD26+ fibroblasts adopt a iCAF signature. These findings may appear to contradict other reports that ascribe a pro-fibrotic phenotype to CD26+ fibroblasts[23,20]. However, our *in vivo* analysis showed that various myCAF-related genes were also upregulated in CD26+ CAFs compared to CD26+ NFs, indicating that CD26+NF-derived iCAFs also possess myofibroblastic functions which may suggest a general activation state inherently related to fibroblasts. In contrast, only a few iCAF-related genes were expressed by CD26-CAFs. Although our functional assays and *in vivo* lineage tracing showed a specific predisposition of NFs towards iCAFs and myCAFs, the *in silico* trajectory analysis did not. The trajectory analysis predicted the transformation of both CD26- and CD26+ NFs into iCAFs first, followed by subsequent transformation into myCAFs. The existence of multiple precursors that give rise to various cellular end-stages complicates these types of *in silico* analyses and may not accurately reflect the biological processes *in vivo.*

CD26+ NFs enhanced tumor cell invasion via MMP-activity and recruited monocytes in a CXCL12-dependent manner, indicating that the CAFs derived from CD26+ NFs have a pro-tumorigenic phenotype. Targeting these CAFs may be especially valuable in the context of ILC, since targeted therapies for this breast cancer subtype are limited and CAFs are abundantly present in these invasive tumors. Further investigation is needed to determine which tumor-secreted factors are important in driving iCAF and myCAF phenotypes and how inhibition of iCAF and myCAF functions may impact tumor progression and therapy response, with the ultimate goal of designing CAF-targeted therapies directed solely at the pro-tumorigenic functions of CAFs.

## Material and Methods

### Reagents

The following flow cytometry antibodies were used throughout this study: EpCAM-PE-Cy7 (Invitrogen, ebioscience clone G8.8), E-cadherin-PE-Cy7 (Biolegend clone DECMA-1), CD49f-AF700 (R&D systems clone GoH3), CD45-AF700 (Invitrogen, ebioscience clone 30-F11), CD45-BUV805 (BD biosciences clone 30-F11), CD45-FITC (Invitrogen, ebioscience clone 30-F11), CD31-BUV395 (BD biosciences clone 390), PDGFRβ-APC (Invitrogen, ebioscience clone APB5), CD26-APC (Biolegend clone H194-112), CD26-PE (Biolegend clone H194-112), CD3-BUV395 (BD biosciences clone 500A2), B220-PE-Cy7 (Invitrogen, ebioscience clone RA3-6B2), CD11b-APC (Invitrogen, ebioscience clone M1/70), CD45-PerCP (BD biosciences clone 30-F11), Sca1-APC-Cy7 (BD biosciences clone D7), CD90.1-FITC (Invitrogen, ebioscience clone HIS51). All flow cytometry antibodies were used in a 1:100 dilution, incubated for 30 minutes on ice. For the whole mount analysis the following antibodies were used: Keratin-14 (rabbit, Covance, PRB155P), alpha-Smooth muscle actin (mouse IgG2a, clone 1A4, Sigma-Aldrich), and CD26 (rabbit monoclonal, Abcam, clone EPR18215). Secondary antibodies: goat anti-rabbit or goat anti-mouse IgG2a conjugated to Alexa-647 (Thermo Fisher, A21245 and A21241 respectively) and donkey anti-rabbit conjugated to Alexa-568 (Thermo Fisher, A10042). The following antibodies were used for immunohistochemistry: PDGFRβ (Cell signaling, #3196), SMA (Thermo scientific, RB-9010), vimentin (Cell signaling, #5741), EpCAM (Abcam, ab32392) and E-cadherin (Cell signaling, #3195). The following neutralizing antibodies were used: anti-CXCL12 (R&D systems, clone 79014, 100 ug/ml) and anti-CXCL2 (Thermo Fisher, clone 40605, 50 ug/ml). The following small molecule inhibitors were used: Sitagliptin (CD26 inhibitor, Tocris, 100 nM), PK-44 (CD26 inhibitor, Tocris, 100 nM), Marimastat (MMP inhibitor, BB-2516, Selleckchem, 100 nM).

### Cell lines

All cells were maintained at 37°C and 5% CO2. WEPtn tumor cell lines were derived from end-stage primary WEPtn tumors. TNBC cell lines were derived from end-stage WB1P primary tumors. Tumor samples were processed into a single cell suspension as described in *‘flow cytometry analysis’* and cultured in DMEM/F12 medium containing 10% FCS, pen/strep (50 units/ml), EGF (5 ng/ml), insulin (5 ug/ml) and cholera toxin (5 ng/ml). Three rounds of differential trypsinization were used to remove fibroblasts from the culture. To ensure all cell lines are depleted from fibroblasts, they were analyzed for their EpCAM expression by flow cytometry. Only cell lines with more than 90% EpCAM+ cells and no expression of PDGFRβ were used for subsequent experiments.

### Mice

Generation of *WapCre;Cdh1^F/F^;Pten^F/F^* (WEPtn) mice has been described previously [25]. *WapCre;Cdh1^F/F^;Col1a1^invCAG-Pik3caH1047R-IRES-Luc^* (WEH1047R) mice were generated by cloning human *Pik3ca* bearing the constitutively activating mutation H1047R in the *Frt-invCag-IRES-Luc* shuttle vector using FseI and PmeI, resulting in *Frt-invCag-Pik3ca^H1047R^-IRES-Luc*. Flp-mediated knockin of the shuttle vector in *WapCre;Cdh1^F/F^* GEMM-ESCs was performed as described previously [53]. Chimeric animals were crossed with *WapCre;Cdh1^F/F^* mice to generate the experimental animal cohorts. Generation of *WapCre;Brca1^F/F^;P53^F/F^* (WB1P) and *WapCre;Brca1^F/F^;Trp53^F/F^;Col1a1^invCAG-Myc-IRES-Luc^* (WB1P-Myc) mice was previously described [54]. *mTmG* reporter mice [55] were backcrossed for seven generations to FVB/n background to accommodate transplantations with donor tissue from our FVB/n-based breast cancer mouse models. *EN1-Cre* mice were purchased from The Jackson Laboratory (JAX stock number:007916) and backcrossed with *mTmG* (FVB/n) mice for 2 generations to generate *EN1-Cre;mTmG* mice for *in vivo* lineage tracing. All mice were housed on standard 12 hr day/night cycle, in individually ventilated cages with ad libitum food. All surgeries were performed under isoflurane anesthesia and rimadyl pain medication. All animal experiments were approved by the Dutch Animal Ethical Committee and conducted in compliance with the Netherlands Cancer Institute and Dutch Animal Welfare guidelines.

### Immunohistochemistry

All immunohistochemical stainings were performed on FFPE material. Slides were deparafinized and rehydrated followed by antigen retrieval in Tris/EDTA (pH 9.0). Endogenous peroxidase was blocked using 3% H2O2. Next the slides were blocked using either 10% non-fat milk in PBS (for EpCAM, PDGFRβ and SMA stainings) or 4% BSA + 5% normal goat serum (NGS) in PBS (for E-cadherin and vimentin stainings). Incubation with primary antibodies was done overnight at 4°C using the following dilutions: PDGFRβ (Cell signaling, #3196) 1:50, SMA (Thermo scientific, RB-9010) 1:200, vimentin (Cell signaling, #5741) 1:200, EpCAM (Abcam, ab32392) 1:200 or E-cadherin (Cell signaling, #3195) 1:200. All primary antibodies were diluted in 1% BSA + 1,25% NGS in PBS. EnVision+ HRP-conjugated anti-rabbit was used as secondary antibody followed by DAB/H202 development and counterstaining with hematoxylin.

### Immunofluorescent labelling and whole-mount imaging

Mammary glands were dissected and incubated in a mixture of collagenase I (1 mg/ml, Roche Diagnostics) and hyaluronidase (50 μg/ml, Sigma Aldrich) at 37 °C for 20 minutes prior to fixation in periodate–lysine–paraformaldehyde (PLP) buffer (1% paraformaldehyde (PFA; Electron Microscopy Science), 0.01M sodium periodate, 0.075 M L-lysine and 0.0375 M P-buffer (0.081 M Na2HPO_4_ and 0.019M NaH2PO_4_; pH 7.4) for 2 h at room temperature. Next, whole glands were incubated in blocking buffer containing 1% bovine serum albumin (Roche Diagnostics), 5% normal goat serum (Monosan) and 0.8% Triton X-100 (Sigma-Aldrich) in PBS for at least 3h at RT. Primary antibodies were diluted in blocking buffer and incubated overnight at room temperature whilst gently shaking. Secondary antibodies diluted in blocking buffer were incubated for at least 8h. Nuclei were stained with DAPI (0.1 μg/ml; Sigma-Aldrich) in PBS. Glands were washed with PBS and mounted on a microscopy slide with Vectashield hard set (H-1400, Vector Laboratories). Primary antibodies: anti-K14 (rabbit, Covance, PRB155P, 1:700), anti-Smooth muscle actin (mouse IgG2a, clone 1A4, Sigma-Aldrich, 1:600), and anti-CD26 (rabbit monoclonal, Abcam, clone EPR18215, 1:200). Secondary antibodies: goat anti-rabbit or goat anti-mouse IgG2a conjugated to Alexa-647 (Thermo Fisher, A21245 and A21241 respectively, 1:400) and donkey anti-rabbit conjugated to Alexa-568 (Thermo Fisher, A10042, 1:400). Whole-mount mammary glands were imaged on an inverted Leica TCS SP8 confocal microscope, equipped with a 405 nm laser, an argon laser, a DPSS 561 nm laser and a HeNe 633 nm laser. Different fluorophores were excited as follows: DAPI at 405 nm, GFP at 488 nm, Tomato or Alexa-568 at 561 nm, and Alexa-647 at 633 nm. Images were acquired with a 25x water immersion objective with a free working distance of 2.40 mm (HC FLUOTAR L 25x/0.95 W VISIR 0.17). Areas of interest were imaged using Z-stacks of 200 μm with an average Z-step size of 2 μm.

### Mouse mammary epithelial cell (MMEC) transplantation

Small tissue fragments of precancerous or control mammary glands were harvested from 4- to 6-week old donor mice and transplanted into the cleared 4^th^ mammary fat pads of 3-week old *mTmG* recipients according to previously published protocols [56,57]. Transplantations in the *En1-Cre;mTmG* recipients were done without clearing of the recipients’ fat pads as it is unclear at which stage and from which direction EN1+ fibroblasts populate the developing mammary gland. For these experiments the donor tissue was placed at the dorsal side of the 4^th^ mammary gland to ensure minimal perturbations to the recipient gland and full potential of all fibroblasts present with the gland. Tumor outgrowth was monitored by palpation and mice were sacrificed at early, advanced and end-stage of tumor growth. Tumors and control tissues were harvested for flow cytometry analysis. For the ILC mouse models (WEPtn and WEH1047R) this required analysis at 18 (early), 24 (advanced) and 30 (end-stage) weeks after transplantation. WB1P and WB1P-Myc derived tumors were analyzed when tumors measured 3×3 mm (early), 8×8 mm (advanced) or 15×15 mm (end-stage).

### Bone marrow transplantation

*Cdh1^F/F^;Pten^F/F^* (EPtn), *Cdh1^F/F^;Col1a1^invCAG-Pik3caH1047R-IRES-Luc^* (EH1047R), *Brca1^F/F^;P53^F/F^* (B1P) and *Trp53^F/F^;Col1a1^invCAG-Myc-IRES-Luc^* (B1P-Myc) recipient mice were lethally irradiated with a single dose of 9 Gy at the age of 8 weeks. Age- and sex-matched *mTmG* donor mice were sacrificed using CO2 and both femurs were isolated for bone marrow harvesting. Femurs were flushed with DMEM + 5% FCS to extract the bone marrow. The cells were filtered over 40 um cell strainers and spun down for 5 minutes at 300 g. The cell pellets were resuspended in PBS and intravenously injected into irradiated mice. Bone marrow from one donor mouse was injected into one recipient mouse. Three weeks after bone marrow engraftment the recipient mice were intraductally injected with lentivirus expressing Cre-recombinase to induce tumor formation. Tumor growth was monitored by palpation and mice were sacrificed at early, advanced and end-stage of tumor growth. For the ILC mouse models (EPtn and EH1047R) this required analysis at 6 (early), 12 (advanced) and 18 (end-stage) weeks after intraductal injection. B1P and B1P-Myc derived tumors were analyzed when tumors measured 3×3 mm (early), 8×8 mm (advanced) or 15×15 mm (end-stage).

### Whole mammary gland transplantation

Whole mammary gland transplantations were performed as described previously by Thompson *et al*.[27] In brief, third mammary glands including nipple and surrounding skin of 4-week old donor mice were isolated and placed in cold PBS while the recipient mouse was prepared for surgery. Recipient mice of 4 weeks old were anaesthetized using isoflurane. A small circular patch of skin was removed between the 3^rd^ and 4^th^ mammary gland, in line with nipples of the recipient mouse. To accommodate the donor gland, the skin was separated from underlying abdominal wall, starting from the incision site dorsally towards the back. A suture was fastened at the dorsal tip of the donor gland and by using the suture the gland was guided in place in the space created along the flank of the recipient mouse. The tip of the donor gland was sutured to the dorsal skin of the mouse. The skin surrounding the nipple of the donor gland was sutured to the skin of the recipient mouse, allowing the ventral side of the donor gland to be positioned on top of the blood vessel running from 3^rd^ to 4^th^ mammary gland. Tumor growth was monitored by palpation and tumors were analyzed at early, advanced and end-stage of tumor growth. For the ILC mouse models (WEPtn and WEH1047R) this required analysis at 12 (early), 18 (advanced) and 24 (end-stage) weeks after transplantation. WB1P and WB1P-Myc derived tumors were analyzed when tumors measured 3×3 mm (early), 8×8 mm (advanced) or 15×15 mm (end-stage).

### Flow cytometry analysis

All tumors and control mammary glands were placed in PBS on ice upon harvesting. Samples were chopped into small pieces using a scalpel and processed into a single cell suspension using a digestion mix containing 2 mg/ml collagenase + 4ug/ml DNase in DMEM/F12. Samples were incubated for 60 minutes at 37°C under continuous shaking. After incubation the collagenase was inactivated by addition of equal volume of DMEM + 5% FCS. Samples were filtered through 70 um cell strainers and spun at 300g for 5 minutes to pellet the cells. Cell pellets were resuspended in red blood cell lysis buffer (RBC lysis, 155 mM NH4Cl, 10 mM KHCO3 and 0.1 mM EDTA in H2O) and incubated on ice for 5 minutes. Next the samples were spun down, 300g for 5 minutes at 4°C. Cell pellets were resuspended in FACS buffer (1% BSA + 5 mM EDTA in PBS) and stained with appropriate antibodies. For WEPtn and WEH1047R the samples were stained with CD45-AlexaFluor700, CD31-BUV395, EpCAM-PE-Cy7, CD26-APC or PDGFRβ-APC. For WB1P and WB1P-Myc the samples were stained with CD45-FITC, CD31-BUV395, EpCAM-PE-Cy7, E-cadherin-PE-Cy7, CD49f-AF700, PDGFRβ-APC or CD26-APC. Tumors transplanted in *En1-Cre*;*mTmG* mice were analyzed using the following antibodies: WEPtn and WEH1047R: EpCAM-PE-Cy7, CD31-BUV395, CD45-BUV805, CD26-APC. WB1P and WB1P-Myc: EpCAM-PE-Cy7, E-cadherin-PE-Cy7, CD49f-AF700, CD31-BUV395, CD45-BUV805. To assess successful engraftment of *mTmG* bone marrow in the bone marrow-transplanted mice we harvested femurs from the mice at time of sacrifice. The femurs were flushed with DMEM + 5% FCS and filtered through a 70 um cell strainer and spun down to pellet cells (300g, 5 minutes). Red blood cells were removed by RBC lysis. Next the samples were spun down (300g, 5 minutes, 4°C) and resuspended in FACS buffer and stained with the following antibodies: CD45-PerCP, Ly6a-APC-Cy7 and CD90.1-FITC. All samples were run on the LSRII SORP flow cytometer with FACS DiVa software version 8.0.1 from Becton Dickinson, San Jose, CA, USA. FlowJo v10 software was used for analysis.

### Sorting and culture of primary fibroblasts

The 3^rd^, 4^th^ and 5^th^ mammary glands were harvested from female mice between 8 and 16 weeks of age. On average 4-6 mice were pooled for one sorting experiment. Samples were processed as described in the section *‘flow cytometry analysis’.* Samples were stained using the following antibodies: EpCAM-PE-Cy7, CD49f-PE-Cy7, CD45-AlexaFluor700, CD31-FITC, CD26-APC. Cells lacking expression of EpCAM, CD49f, CD45 and CD31 were considered fibroblasts. Sorting was done on a BD FACS Aria Fusion at 20psi using a 100um nozzle. CD26- and CD26+ fibroblasts were collected, spun down (300 g, 5 minutes) and plated in collagen type I-coated plates (8 ug/cm^2^) in DMEM + 20% FCS. All primary fibroblasts were cultured for no more than 6 passages. Experiments using primary fibroblasts were performed at the lowest possible passage numbers, typically passage 2 to 3.

### Transwell assay

All fibroblast recruiting transwell assays were performed in 24-well set-up using Corning inserts (8.0 um pores, transparent PET membrane, 1×10^5^ pores per cm^2^) and companion plates. Inserts were coated with growth-factor reduced matrigel (Corning) diluted in serum-free medium (DMEM) to a concentration of 0.25 mg/ml protein. Tumor cells or conditioned medium derived from tumor cells (1×10^6^ cells per 10 cm dish in serum-free medium for 24 hours) were placed in the bottom wells. In each well 1×10^5^ tumor cells were plated. 24 hours after plating the culture medium was replaced by serum-free medium (DMEM/F12). Coated inserts were placed in bottom wells and 5×10^4^ fibroblasts were plated in the inserts in DMEM + 0.2% FCS. After 24 hours the inserts were harvested, cleared of cells remaining in the top compartment using cotton swaps and fixed in ice-cold methanol for 5 minutes. Next the inserts were stained with crystal violet (0.5% in 2:1 H2O:methanol) for 5 minutes. Inserts were cleaned in tap water and left to dry prior to imaging.

### Organotypic invasion assay

CD26- or CD26+ NFs were plated in 1 ml gel composed of collagen (5 mg/ml):BME:medium (2:1:1) in 24-well suspension plates. 4×10^5^ fibroblasts were plated in each gel and left to solidify at 37°C, 5% CO2 for 1 hour. Next tumor cells were plated on fibroblast-containing or empty gels at a density of 1×10^5^ cells. One day after plating, the tumor cells were covered with 100 ul BME:medium (1:1) to prevent disruption of the tumor cells monolayer during transfer. One hour after sealing the tumor cells, the gels were lifted from the plate and transferred to stainless-steel grids in a 6-well plate, allowing the gels to be surrounded by medium. The gels were cultured for 1 week in a 1:1 mix of tumor cell culture medium and fibroblast culture medium (hereafter referred to as mixed medium) in the absence or presence of Marimastat (100 nm) or neutralizing antibodies against CXCL12 (100 ug/ml). Medium was changed after 3 days. One week after plating the gels were harvested and fixed in 4% formalin and processed to FFPE slides for HE and EpCAM staining.

### Splenocyte recruitment assay

Conditioned medium (CM) was derived from mono or co-cultures of CD26- NFs, CD26+ NFs or tumor cells. CD26- or CD26+ NFs were plated in a 6-well plate at a density of 2×10^5^ cells. Tumor cells (WEPtn derived) were plated in a 6-well plate at a density of 5×10^4^ cells. For the co-cultures 2×10^5^ fibroblasts were plated together with 5×10^4^ tumor cells. All conditions were cultured in mixed medium for 3 days. Spleens were harvested from non-tumor bearing mice between 10 and 16 weeks of age and chopped into small pieces and digested briefly (15 min) in digestion mix, filtered and cleared of red blood cells (see section *‘flow cytometry analysis’).* The splenocytes were resuspended in mixed medium and counted. All splenocyte recruitment assays were performed in 6-well set-up using Corning inserts (8.0 um pores, transparent PET membrane, 1×10^5^ pores per cm^2^) and companion plates. Inserts were coated with growth-factor reduced matrigel (Corning) diluted in mixed medium to a concentration of 0,5 mg/ml protein. Coated inserts were placed in bottom wells with CM and 5×10^5^ splenocytes were plated in the inserts. After 24 hours the inserts were carefully removed and the bottom well contents were collected and spun down (300g, 5 min). Pellets were resuspended in FACS buffer and stained using the following antibodies: CD11b-APC, CD3-BUV395 and B220-PE-Cy7. All samples were analyzed on the LSRII from BD biosciences. FlowJo v10 software was used for analysis.

### Cytokine array

WEPtn-derived tumor cells (13-MCB-17) were co-cultured with either CD26- NFs or CD26+ NFs at a density of 2,5×10^5^ tumor cells with 1×10^6^ fibroblasts in a 10 cm dish in mixed medium for three days. Conditioned medium was harvested and spun at 2000 g for 5 minutes at 4°C to pellet any cells or debris. The supernatant was used for the cytokine array (Proteome Profiler Mouse Cytokine Array Kit, Panel A, R&D systems) according to the manufacturer’s protocol.

### Gene expression analysis

Based on the amount of cells homogenized in the RLT buffer (79216, Qiagen), the total RNA was isolated using the RNeasy Mini Kit (74106, Qiagen) and RNeasy Micro Kit (74004, Qiagen), including an on column DNase digestion (79254, Qiagen), according to the manufacturer’s instructions. Quality and quantity of the total RNA was assessed on the 2100 Bioanalyzer instrument following manufacturer’s instructions “Agilent RNA 6000 Nano” (G2938-90034, Agilent Technologies) and “Agilent RNA 6000 Pico” (G2938-90046, Agilent Technologies). Total RNA samples having RIN>7 were subjected to TruSeq stranded mRNA library preparation, according to the manufacturer’s instructions (Document # 1000000040498 v00, Illumina). The stranded mRNA libraries were analyzed on a 2100 Bioanalyzer instrument following the manufacturer’s protocol “Agilent DNA 7500 kit” (G2938-90024, Agilent Technologies), diluted to 10nM and pooled equimolar into multiplex sequencing pools for sequencing on the HiSeq 2500 and NovaSeq 6000 Illumina sequencing platforms. HiSeq 2500 single-end sequencing was performed using 65 cycles for Read 1, 10 cycles for Read i7, using HiSeq SR Cluster Kit v4 cBot (GD-401-4001, Illumina) and HiSeq SBS Kit V4 50 cycle kit (FC-401-4002, Illumina). NovaSeq 6000 paired-end sequencing was performed using 54 cycles for Read 1, 19 cycles for Read i7, 10 cycles for Read i5 and 54 cycles for Read 2, using the NovaSeq6000 SP Reagent Kit v1.5 (100 cycles) (20028401, Illumina). Differential gene expression was determined using the R-package DESeq2[58] with the following criteria: FDR-corrected p-value <0.05 and log2FoldChange>2 (up-regulated) or log2FoldChange<-2 (down-regulated). Hierarchical clustering was performed using Euclidean distance. For the pathway analyses a Fisher’s exact test was used with the KEGG or GO geneset from MSigDB. Single sample gene set enrichment was calculated with the R-package Singscore[59].

### Single-cell transcriptomics on fibroblasts from WEPtn mice

*Cdh1^F/F^;Pten^F/F^* mice (EPtn, 7-8 weeks of age) were intraductally injected with lentivirus expressing Cre-recombinase to induce tumor formation or PBS as controls. Cre-injected glands were harvested 6, 12 or 18 weeks after injected. PBS-injected glands were harvested 12 weeks after injection. Glands/tumors were processed into a single cell suspension as described in the section *‘flow cytometry analysis’.* Samples were stained using the following antibodies: CD31-APC, CD45-AF700, EpCAM-FITC and CD49f-PE-Cy7. Fibroblasts were considered negative for all mentioned markers. Sorted fibroblasts were spun down and frozen in FCS + 10% DMSO until use. On day of scRNA sequencing, samples were thawed, checked for viability by flow cytometry (percentage of dapi-negative cells was above 75% for all samples). Cells were resuspended 1000 cells/ul in 1xPBS containing 0.04% weight/volume BSA (400 ug/ml) and for each sample the Chromium Controller platform of 10X Genomics was used for single cell partitioning and barcoding. Per sample each cell’s transcriptome was barcoded during reverse transcription, pooled cDNA was amplified and Single Cell 3’ Gene Expression libraries were prepared according to the manufacturer’s protocol “Chromium Next GEM Single Cell 3’ Reagent Kits v3.1” (CG000204, 10X Genomics). All four Single Cell 3’ Gene Expression libraries were quantified on a 2100 Bioanalyzer Instrument following the manufacturer’s protocol “Agilent DNA 7500 kit” (G2938-90024, Agilent Technologies). These Single Cell 3’ Gene Expression libraries were combined to create one sequence library pool which was quantified by qPCR, according to manufacturer’s protocol “KAPA Library Quantification Kit Illumina® Platforms” (KR0405, KAPA Biosystems). The NextSeq 550 Illumina sequencing system was used for paired end sequencing of the Single Cell 3’ Gene Expression libraries at a sequencing depth of approximately 18.000 reads pairs/cell. NextSeq 550 paired end sequencing was performed using 28 cycles for Read 1, 8 cycles for Read i7 and 56 cycles for Read 2, using NextSeq 500/550 High Output Kit v2.5 (75 Cycles) Reagent Kit (PN 20024906, Illumina). The data was analyzed using the Python package Scanpy. Cells containing less than 200 genes and genes present in less than 3 cells were excluded from the data. Remaining cells that contained expression of the marker genes (*Cd31, Cd45, Epcam, Cd49f)* used for sorting were excluded from the data. Additional filtering and cutoffs were placed based on distribution plots. Cells with gene counts between 1000 and 4000 and with a mitochondrial percentage of less than 20% were included in analysis. Clustering was made using the Leiden algorithm. Trajectory analysis was performed using the PAGA[60].

### Single-cell transcriptomics on WB1P and WB1P-Myc tumors

End-stage WB1P and WB1P-Myc tumors were digested into a single cell suspension as described in the section *‘Flow cytometry* analysis’. Single cell suspensions of the tumors were sorted by FACS to isolate live single cells (FSC-A x FSC-H to define singlets, dapi-negative cells to define live cells). After sorting the cells were subjected to Dropseq RNA sequencing according to the protocol of Macosko *et al*.[61] using WB1P cells (275 cells/ul) and WB1P-Myc cells (270 cells/ul) both in 1xPBS+0.01%BSA. Dropseq beads ‘MACOSKO-2011-10’ were purchased from ChemGenes. Ready-made Drop-Seq microfluidic devices were purchased from Nanoshift. During droplet generation, of both samples multiple 5 minute fractions of droplets were collected and based on droplet quality assessment for each sample 4 fractions were selected for further processing according to “Drop-seq laboratory protocol v3.1”. After breakage, reverse transcription and Exocuclease I treatment, of each fraction the cDNA was amplified in triplicate PCR reactions. After PCR the amplified cDNA products were pooled, cleaned by a 0.6X AMPure XP bead cleanup and quantified on a 2100 Bioanalyzer Instrument following the manufacturer’s protocol “Agilent High Sensitivity DNA Kit” (G2938-90321, Agilent Technologies). This procedure was done twice for every fraction selected. The amplified cDNA product was concentrated by speedvac and all was used as input for the final Nextera XT library preparations. All Libraries were quantified on a 2100 Bioanalyzer Instrument following the manufacturer’s protocol “Agilent High Sensitivity DNA Kit” and diluted to 10 nM before paired-end sequencing on the MiSeq and HiSeq2500 Illumina sequencing platforms. Sequencing was performed using 25 cycles for Read 1, 8 cycles for Read i7 and 117 cycles for Read 2, using MiSeq Reagent Kit v3 (150-cycle) (MS-102-3001, Illumina), HiSeq PE Rapid Cluster Kit v2 (PE-402-4002, Ilumina) and HiSeq Rapid SBS Kit v2 (200 cycles) (FC-402-4021, Illumina). Sequencing was performed on the MiSeq and HiSeq2500 Illumina sequencing platforms. The data was analyzed using the Python package Scanpy. Cells containing less than 200 genes and genes present in less than 3 cells were excluded from the data. Additional filtering and cutoffs were placed based on distribution plots. Cells with gene counts between 450 and 4000 and with a mitochondrial percentage of less than 10% were included in analysis.

## Statistical analysis

The statistical analyses used throughout the manuscript have been indicated in the figure legends.

## Data availability

The raw data generated in this study is available upon request from the corresponding authors.

## Author contributions

Julia M. Houthuijzen performed and designed the experiments and wrote the paper together with Jos Jonkers. Roebi de Bruijn performed all bioinformatic analyses on the single-cell transcriptomics datasets and other *in vitro* and *in vivo* RNA sequencing experiments. Eline van der Burg and Anne-Paulien Drenth assisted with the animal experiments and performed tumor volumetric measurements on all mice. Ellen Wientjens was responsible for genotyping of the mice used within these experiments and assisted with *in vitro* experiments. Tamara Filipovic assisted with the analysis of tumors from the transplantations in the En1-Cre;*mTmG* mice. Esme Bullock performed the iCAF and myCAF signature analyses on the human ILC and IDC laser microdissected tumor specimens. Chiara S. Brambillasca prepared the samples for single-cell transcriptomics of the WB1P and WB1P-Myc tumors. Marja Nieuwland and Iris de Rink performed the single-cell transcriptomics and quality control of RNAseq data. Frank van Diepen assisted with FACS sorting of tumor samples and co-cultures for sequencing. Sjoerd Klarenbeek generated the WEPtn mouse model. Ron Kerkhoven was head of sequence facility and assisted in the design of single-cell transcriptomics experiments. Valerie Brunton published and provided the human ILC and IDC datasets. Colinda Scheele performed whole mount analyses on EN1-Cre;*mTmG* transplanted mice and wildtype mice. Mirjam C. Boelens initiated the CAF research in WEPtn mice. Jos Jonkers is head of the Molecular Pathology department and wrote the paper together with Julia Houthuijzen.

## Acknowledgements

We would like to thank the people at the Genomics Core Facility of the Netherlands Cancer Institute, especially Wim Brugman, for all the RNA isolations, RNA library preps and sequence runs presented in this study.

