## Supplemental figures for "CD26-negative and CD26-positive tissue-resident fibroblasts contribute to functionally distinct CAF subpopulations in breast cancer"

### Supplemental figure 1

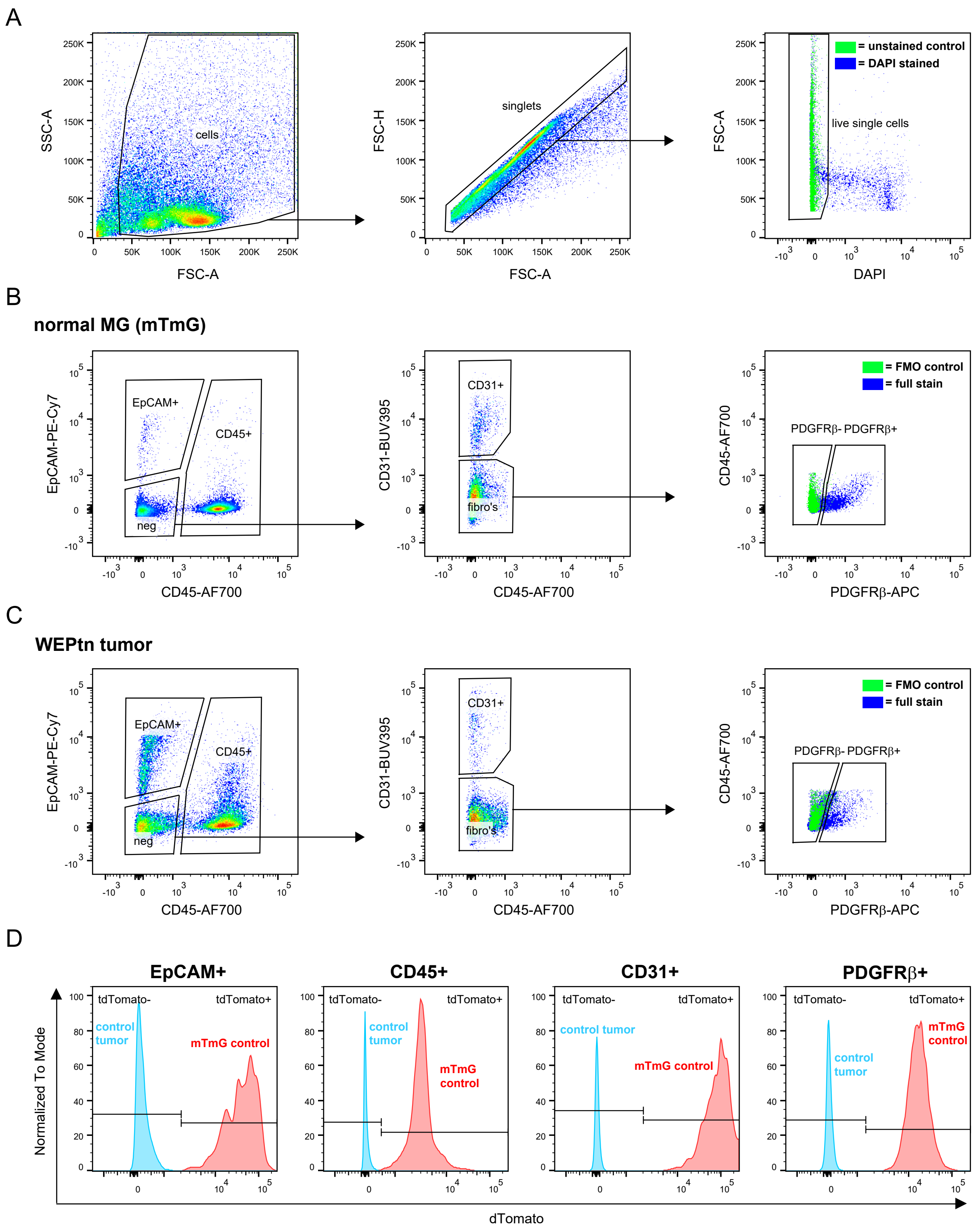

Supplemental figure 2

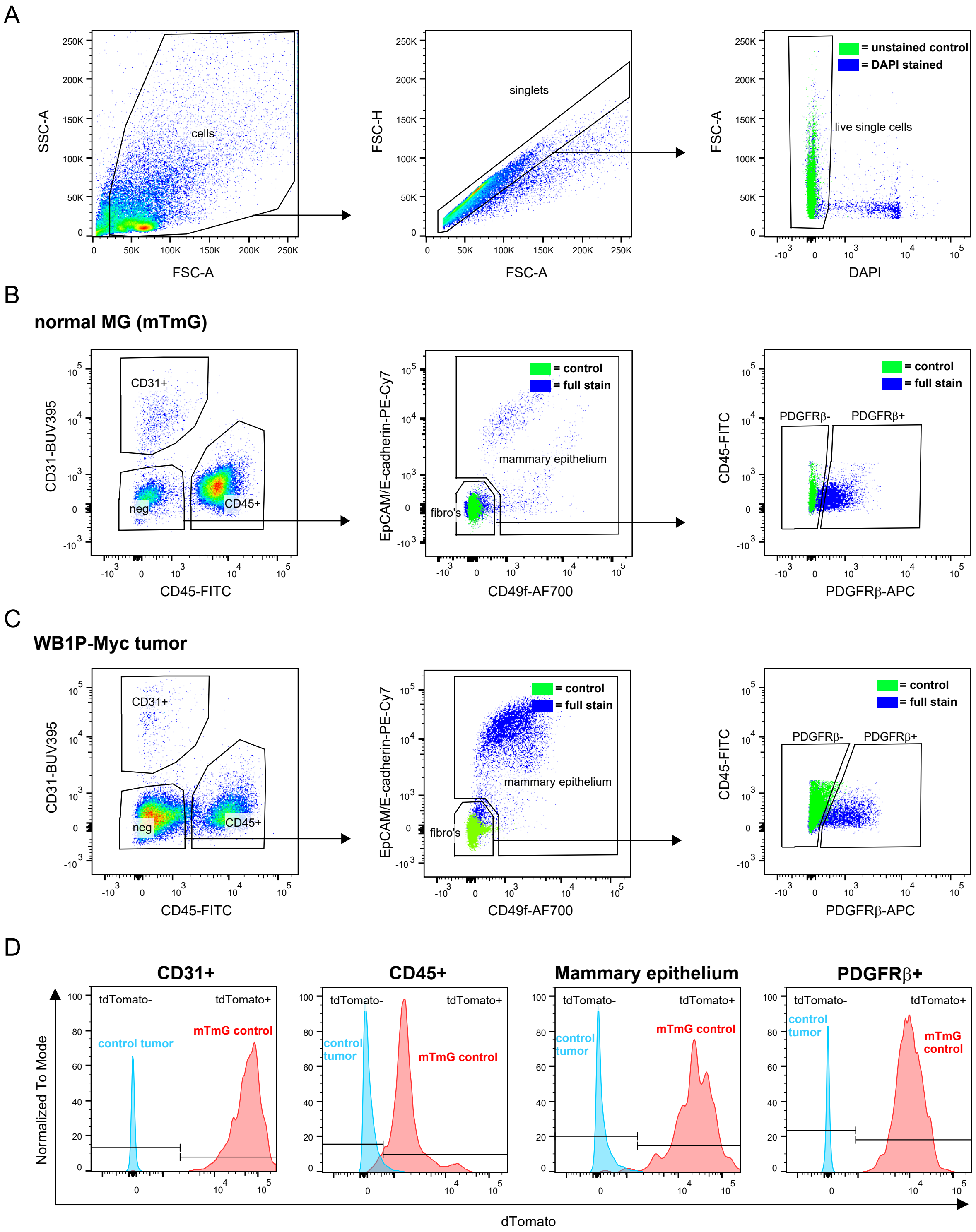

Supplemental figure 3

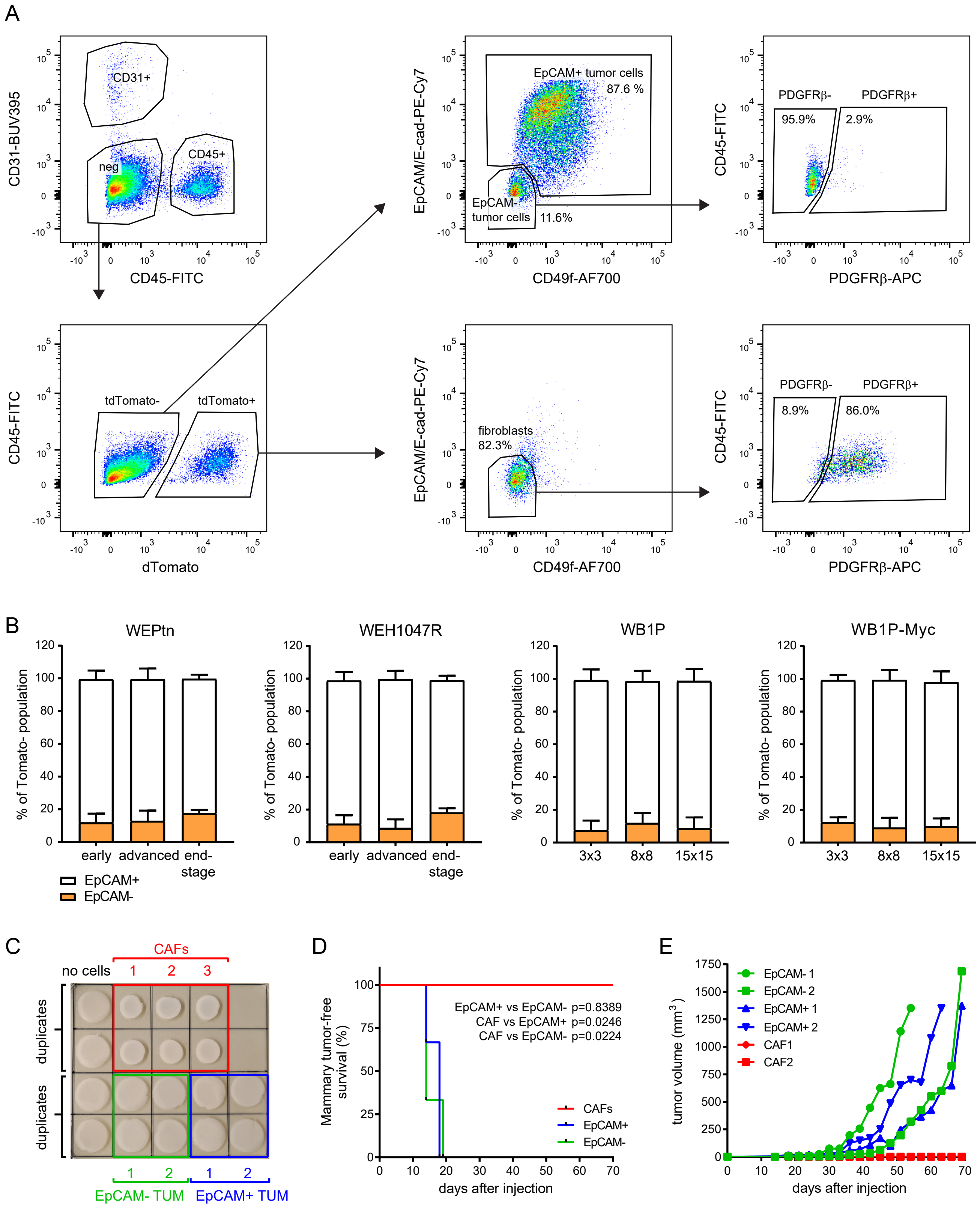

Supplemental figure 4

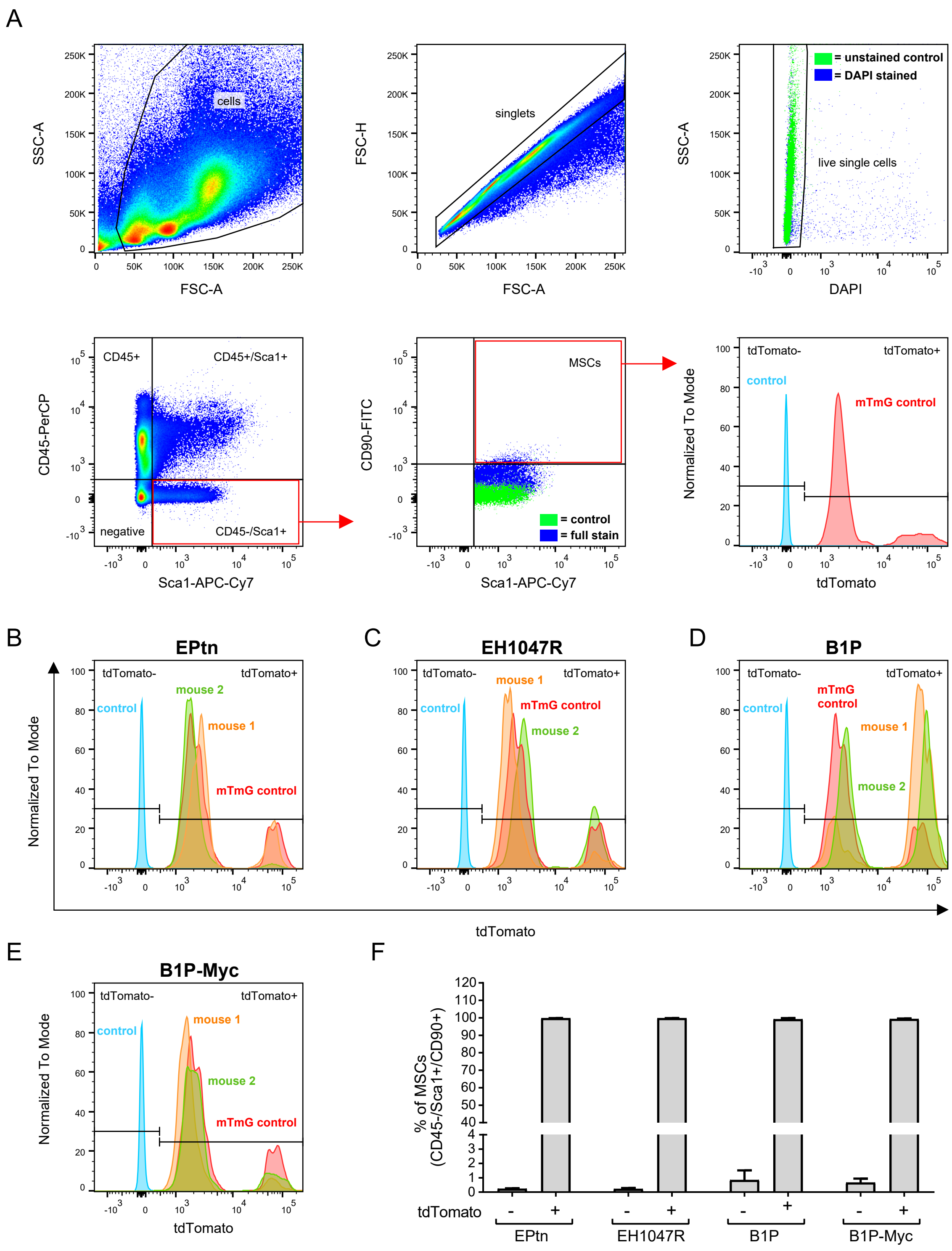

### Supplemental Figure 5

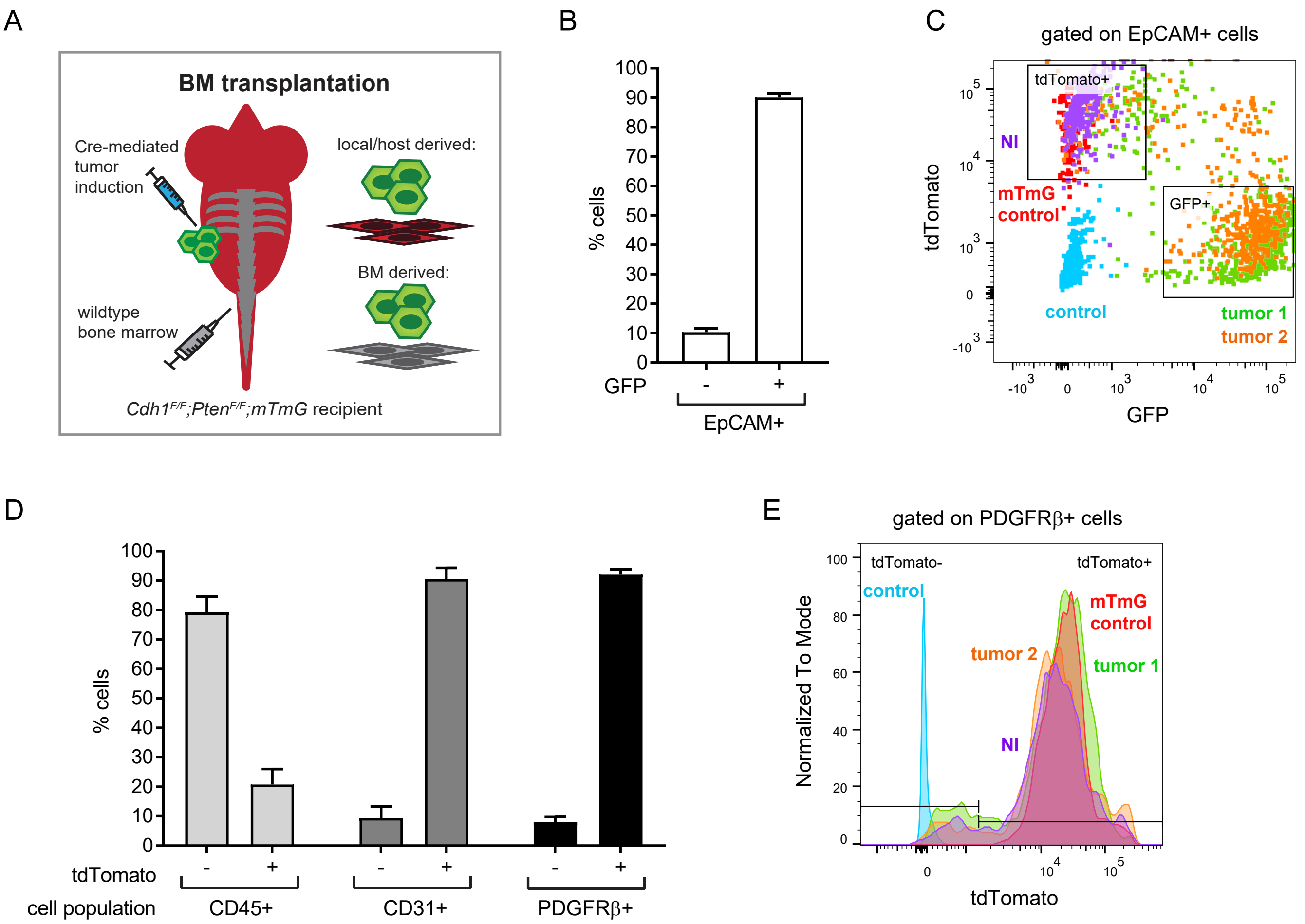

Supplemental figure 6

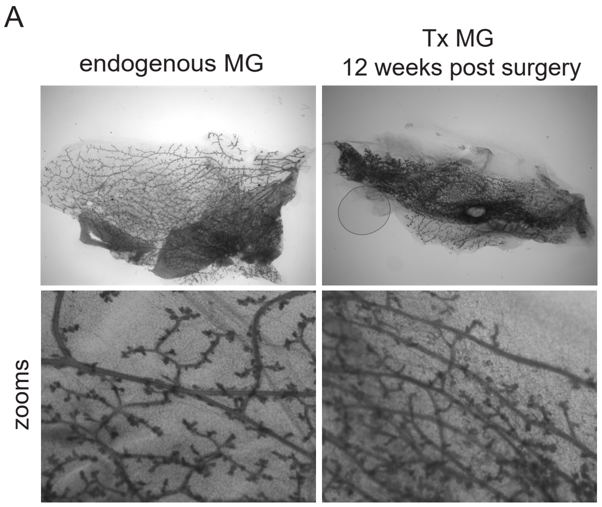


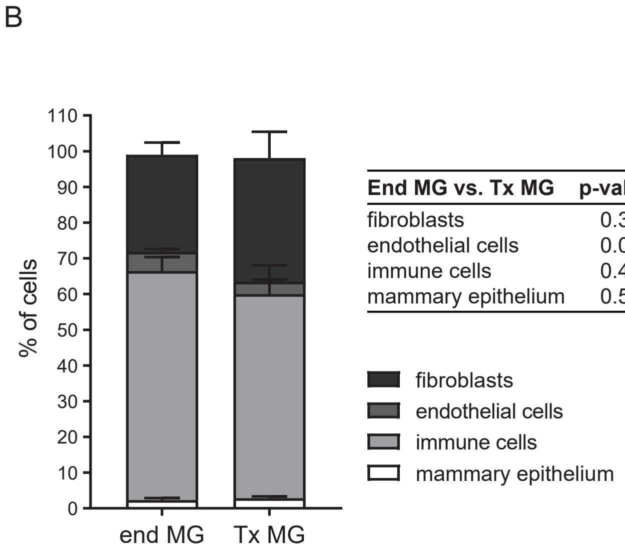


| **End MG vs. Tx MG** | **p-value** |
|---|---|
| fibroblasts | 0.388 |
| endothelial cells | 0.073 |
| immune cells | 0.458 |
| mammary epithelium | 0.507 |

### Supplemental figure 7

A

**ILC iCAF signature**

| | | |
|---|---|---|
| *C4b* | *Cfb* | *Fgf7* |
| *C3* | *Mmp3* | *Serping1* |
| *Serpina3n* | *Bgn* | *Apod* |
| *Ly6a* | *Ccn5* | *Steap4* |
| *Gas6* | *Ccl8* | *Txn1* |
| *Igf1* | *Lbp* | *Il1r1* |
| *Cxcl12* | *Pcolce* | *Ccl7* |
| *Il11ra1* | *Ly6c1* | |
| *Tmem176b* | *Tmem176a* | |

**ILC myCAF signature**

| | | |
|---|---|---|
| *Pmepa1* | *Serpine2* | *Mfap4* |
| *Bgn* | *Cilp* | *Cdh11* |
| *Olfml3* | *Mgp* | *Tnc* |
| *Spon1* | *Timp1* | *Bpgm* |
| *Col12a1* | *Ccn4* | *Igfbp3* |
| *Runx1* | *Itgb5* | *Fzd1* |
| *Lrrc15* | *Postn* | *F2r* |
| *Cpxm2* | *Ecrg4* | |
| *Col8a1* | *Gpc1* | |

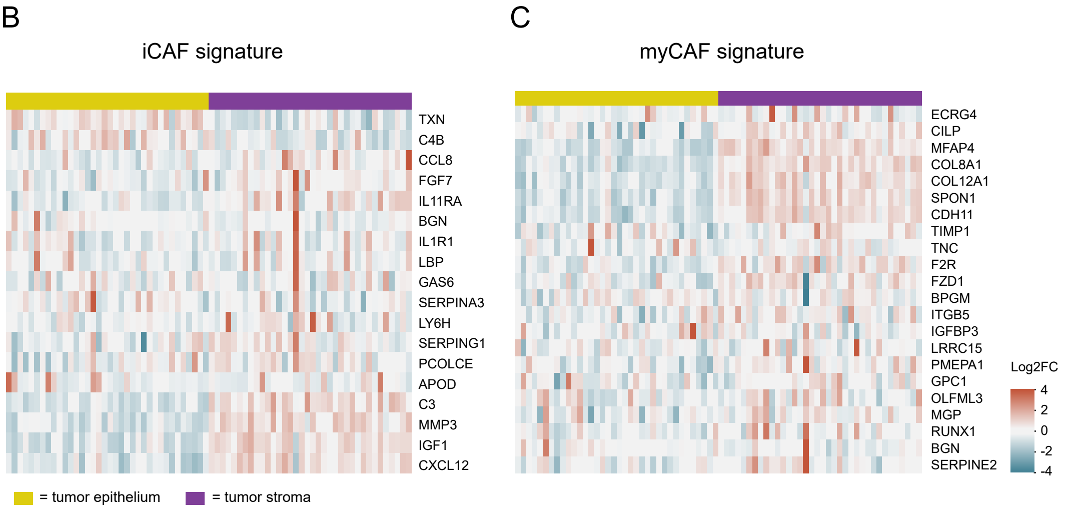


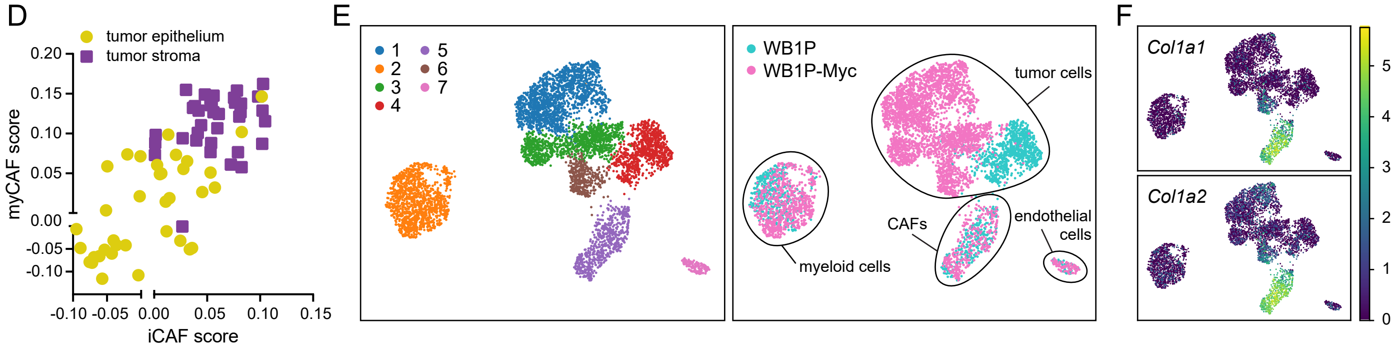

Supplemental figure 8

A

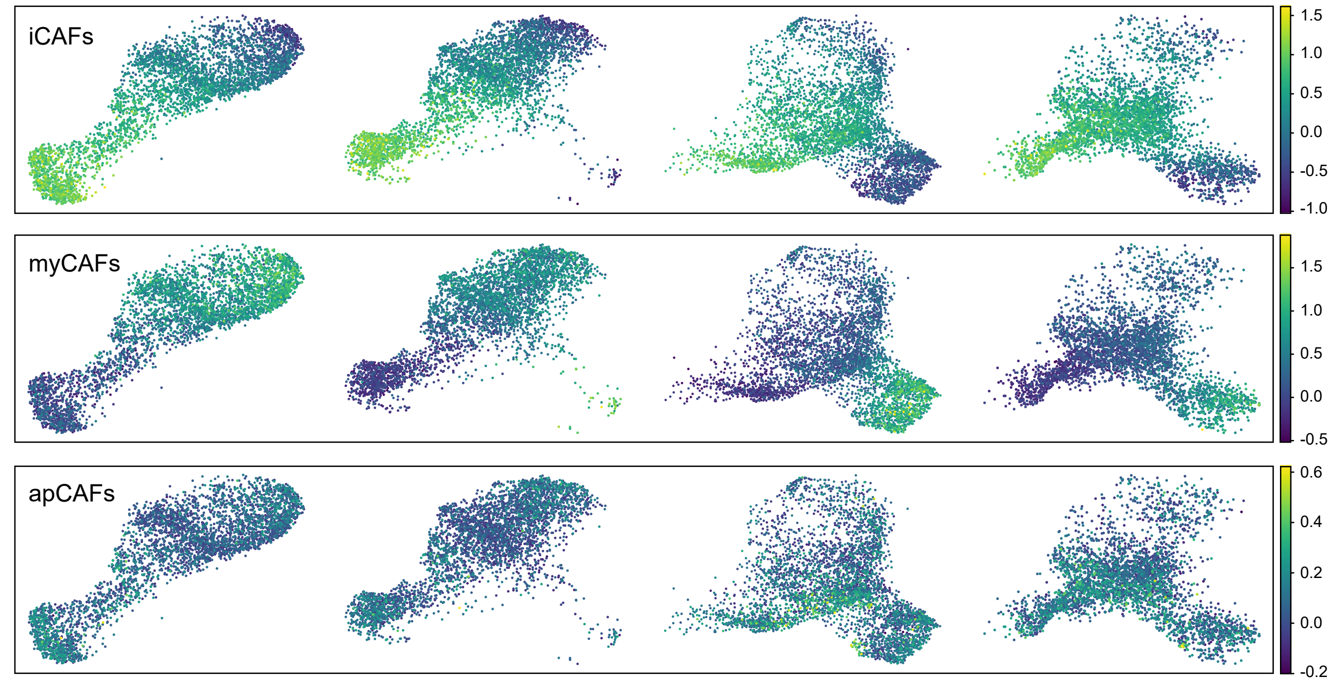


B

| **PDAC iCAF signature** | | |
|---|---|---|
| *Dpt* | *Scara3* | *Ly6a* |
| *Ifi205* | *Pcolce2* | *Ly6c1* |
| *Gsn* | *Clec3b* | *Adamts5* |
| *Fifg* | *Efemp1* | *Tnxb* |
| *Svep1* | *Ifi27l2a* | *C4b* |
| *Plpp3* | *Col14a1* | *C3* |

| **PDAC myCAF signature** | | |
|---|---|---|
| *Serpine2* | *Crlf1* | *Sdc1* |
| *Col15a1* | *Thy1* | *Cthrc1* |
| *Tnc* | *Tagln* | *Col8a1* |
| *Sparcl1* | *Col12a1* | *Thbs2* |
| *Spp1* | *Col1a1* | *Acta2* |
| *Tgfb1* | *Cxcl14* | |

| **PDAC apCAF signature** | | |
|---|---|---|
| *Slpi* | *Lgals7* | *Krt8* |
| *Ptgis* | *Spint2* | *Cxadr* |
| *Nkain4* | *Dmkn* | *Ezr* |
| *Hspb1* | *Saa3* | *H2-Ab1* |
| *Cav1* | *Clu* | *Cd74* |
| *Arhgdib* | *Slc9a3r1* | *Fth1* |

C D

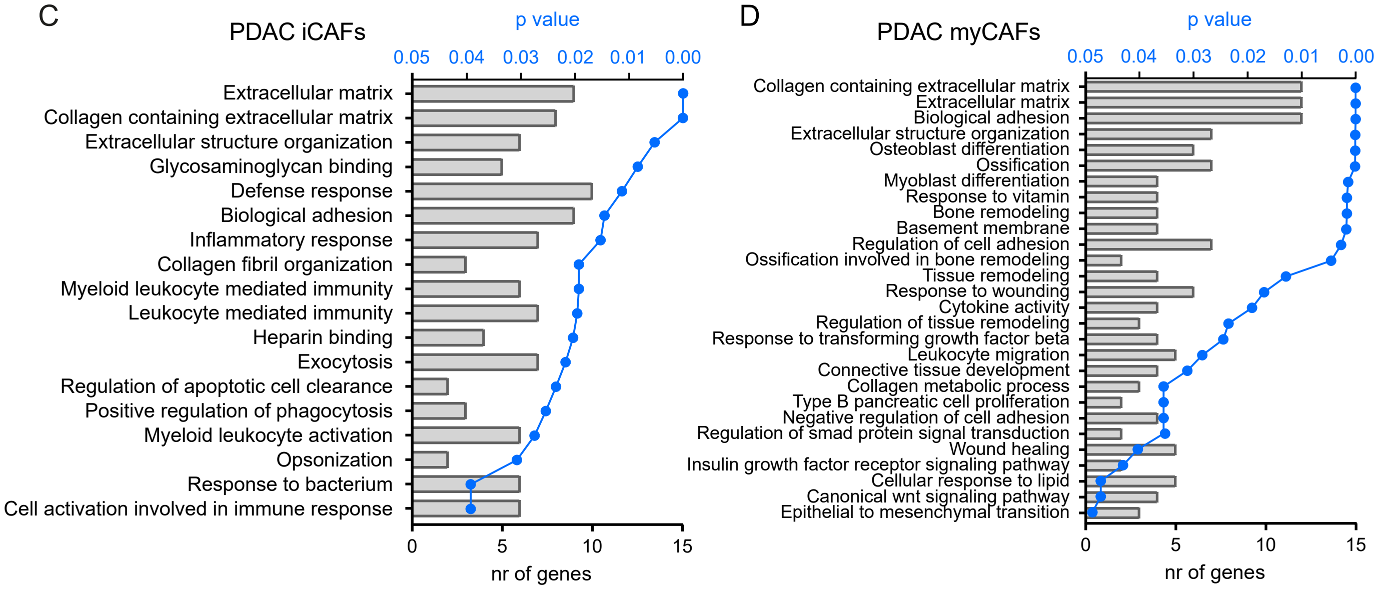


E F

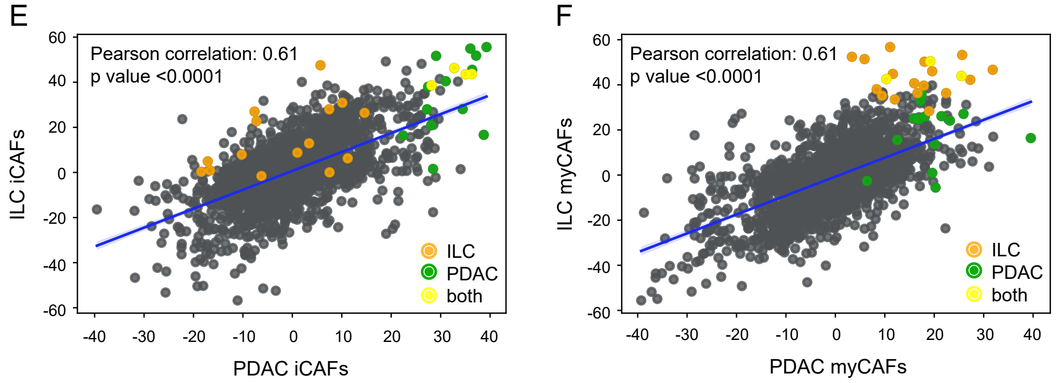

Supplemental figure 9

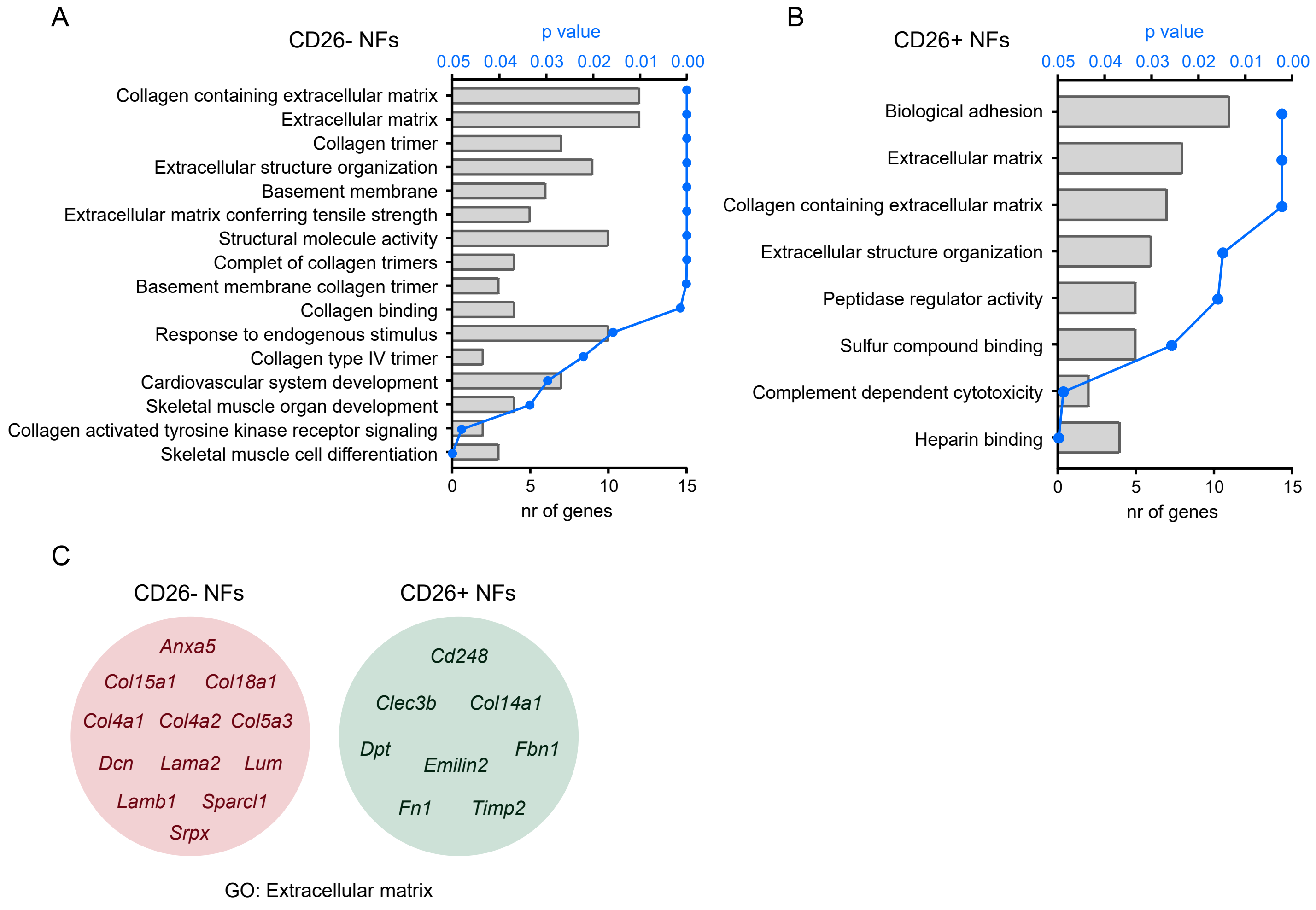

Supplemental figure 10

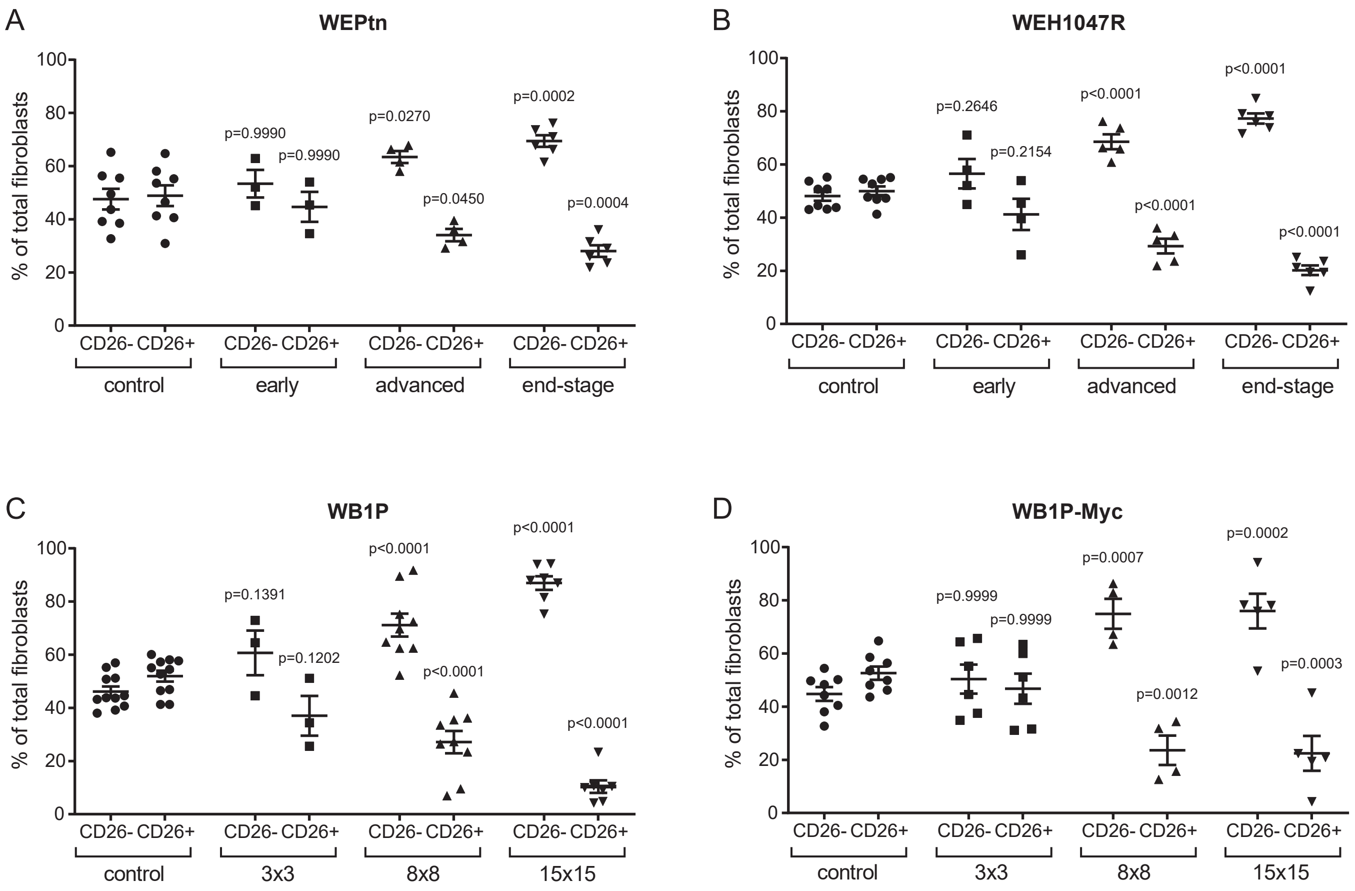

Supplemental figure 11

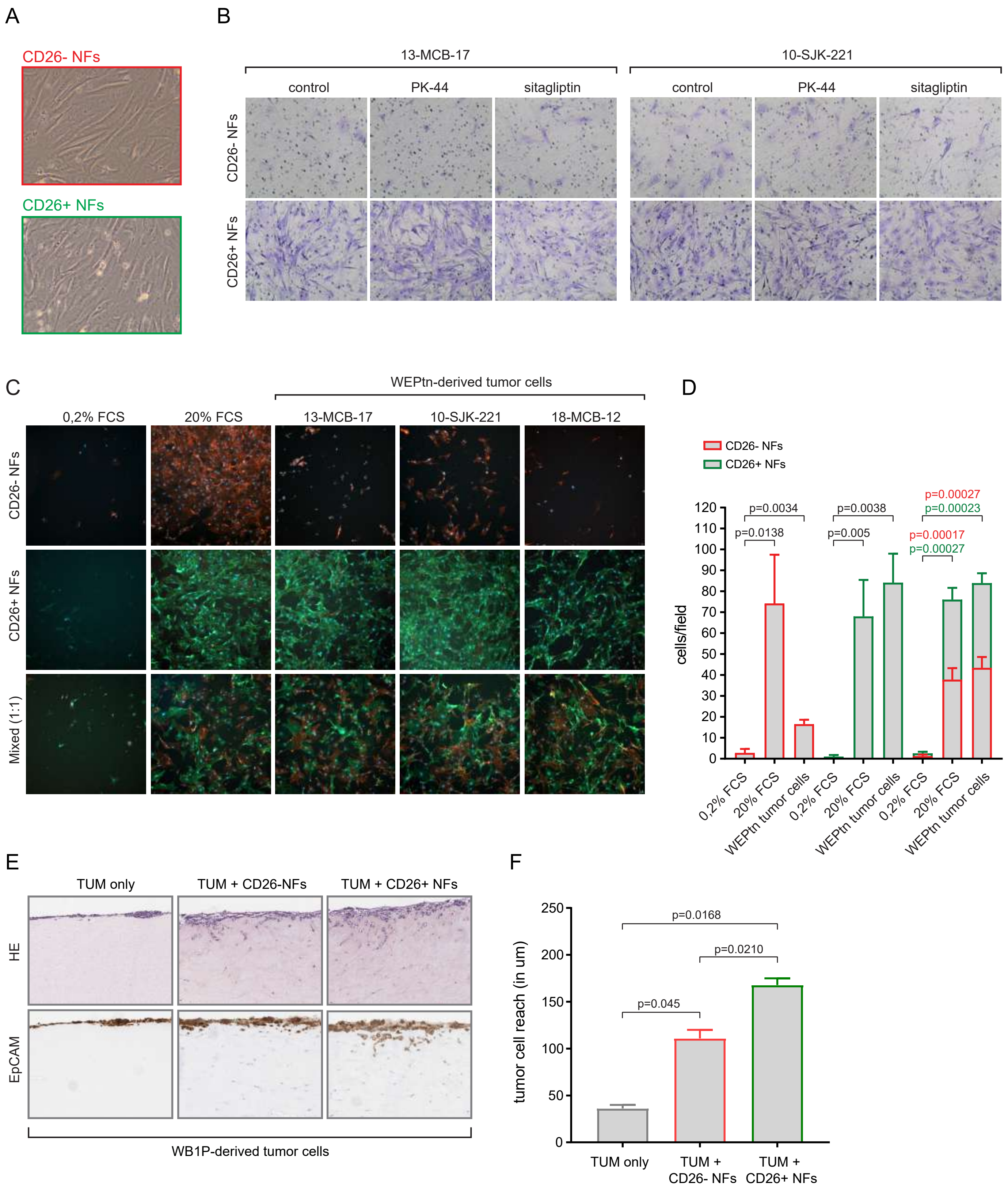

#### Supplemental figure 12

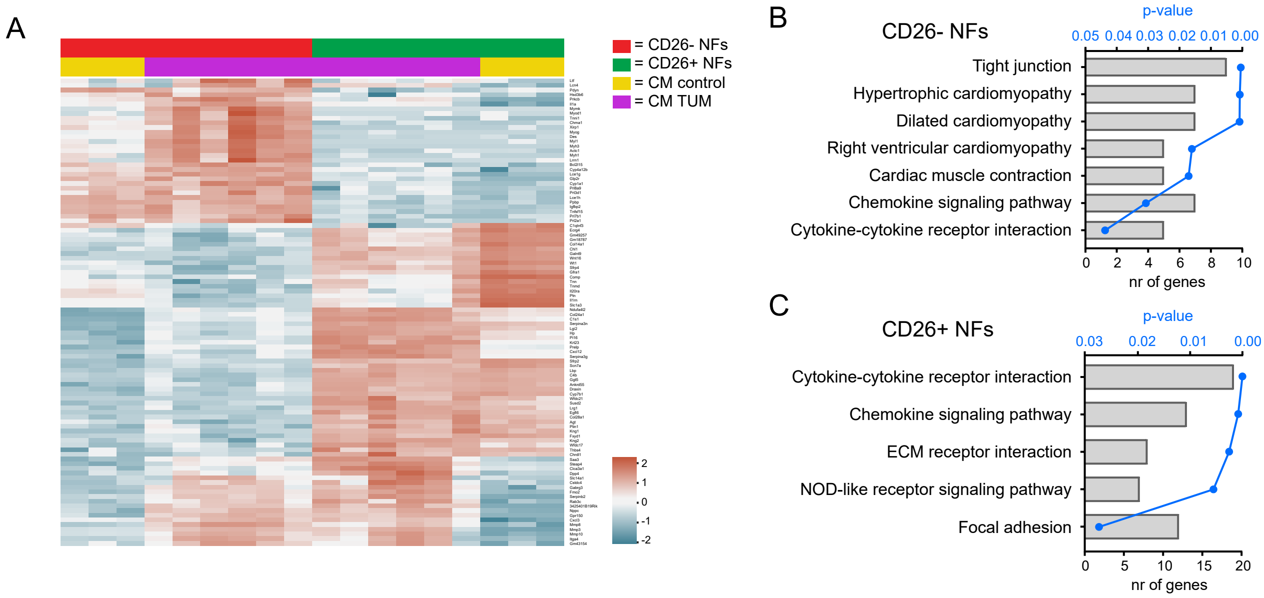

Supplemental figure 13

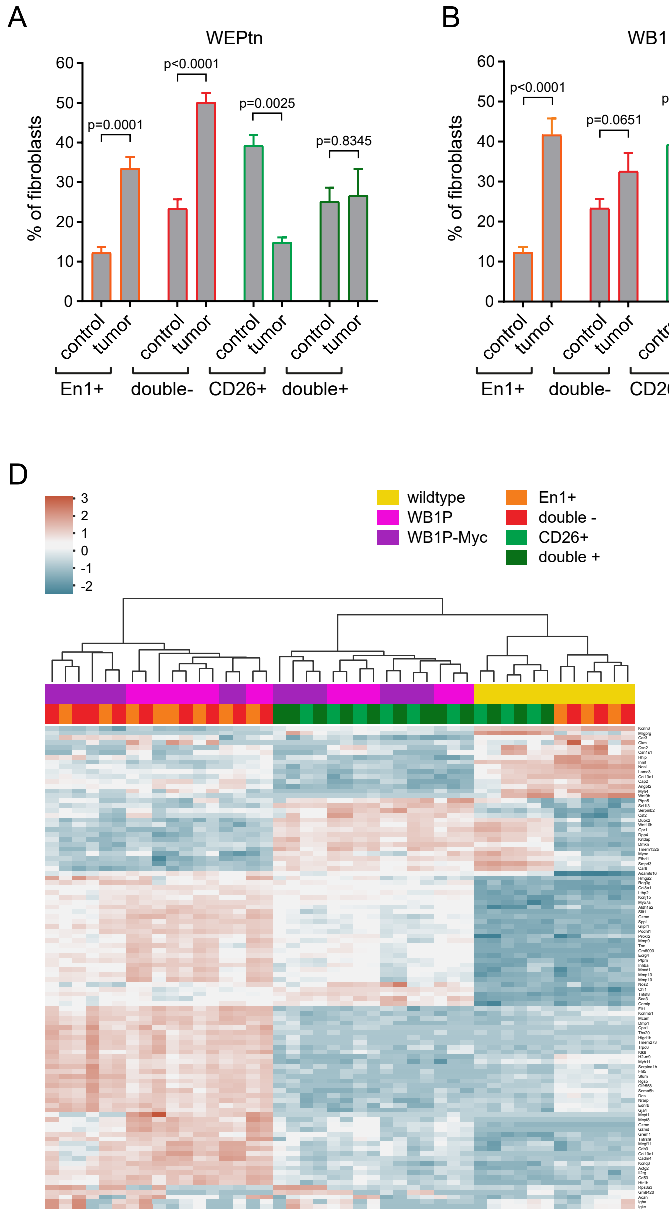

Supplemental figure 14

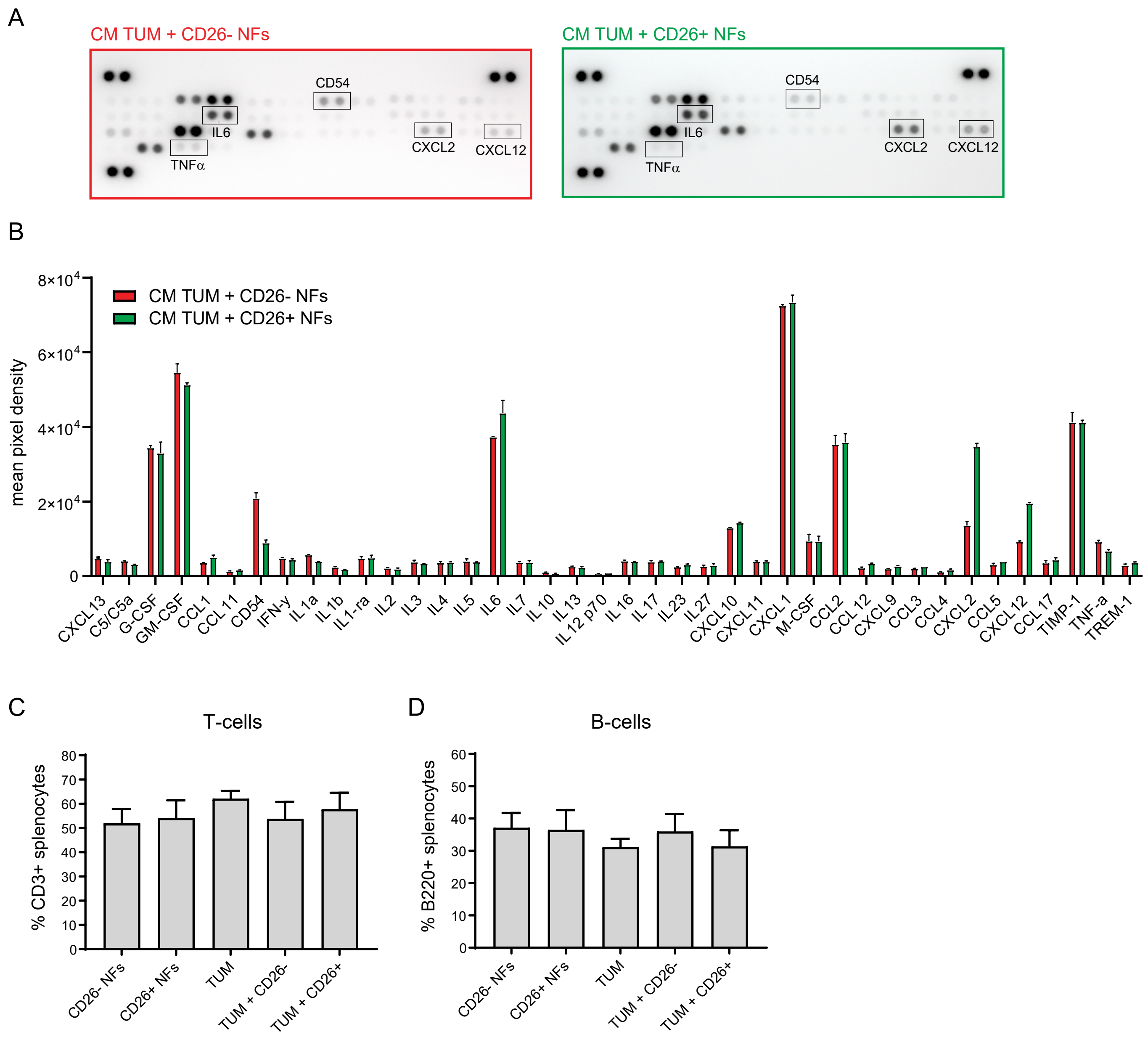
