## Supplemental figure legends for "CD26-negative and CD26-positive tissue-resident fibroblasts contribute to functionally distinct CAF subpopulations in breast cancer"

**Supplemental figure 1: Gating strategy for WEPtn and WEH1047R tumors and controls from transplantation studies.** A) General gating done on all samples for cells, singlets and live single cells. B) Normal mammary glands derived from mTmG mice were stained with EpCAM-PE-Cy7, CD45-AF700, CD31-BUV395 and PDGFRβ-APC. Mammary epithelium was defined as EpCAM+/CD45-/CD31-/PDGFRβ-. Immune cells were defined as CD45+/CD31-/EpCAM-. Endothelial cells were defined as CD31+/EpCAM-/CD45-. Fibroblasts were defined as EpCAM-/CD45-/CD31-/PDGFRβ+. C) WEPtn and WEH1047R derived ILC tumors (WEPtn shown here) were stained with EpCAM-PE-Cy7, CD45-AF700, CD31-BUV395 and PDGFRβ-APC and used the same markers to identify epithelium, immune cells, endothelium and fibroblasts as in panel B. In panel B and C fluorescence-minus-one (FMO) control stained samples (all antibodies except PDGFRβ) are shown in green and full stained samples are shown in blue. D) tdTomato expression in indicated cell populations from mTmG control mice and WEPtn mice with gates that define tdTomato+ (in red) and tdTomato- (in blue) cells for each cell population.

**Supplemental figure 2: Gating strategy for WB1P and WB1P-Myc tumors and controls from transplantation studies.** A) General gating done on all samples for cells, singlets and live single cells. B) Normal mammary glands derived from mTmG mice were stained with CD31-BUV395, CD45-FITC, EpCAM-PE-Cy7, E-cadherin-PE-Cy7, CD49f-AF700 and PDGFRβ-APC. Mammary epithelium was defined as EpCAM/E-Cadherin+/CD49f+/low/CD45-/CD31-/PDGFRβ-. Immune cells were defined as CD45+/CD31-/EpCAM/E-Cadherin-. Endothelial cells were defined as CD31+/EpCAM/E-Cadherin-/CD45-. Fibroblasts were defined as EpCAM/E-Cadherin-/CD49f-/CD45-/CD31-/PDGFRβ+. C) WB1P and WB1P-Myc derived TNBC tumors (WB1P-Myc shown here) were stained with CD31-BUV395, CD45-FITC, EpCAM-PE-Cy7, E-cadherin-PE-Cy7, CD49f-AF700 and PDGFRβ-APC and used the same markers to identify epithelium, immune cells, endothelium and fibroblasts as in panel B. In panel B and C control stained samples (only CD31-BUV395 and CD45-FITC) are shown in green and full stained samples are shown in blue. D) tdTomato expression in indicated cell populations from mTmG control mice and WB1P-Myc mice with gates that define tdTomato+ (in red) and tdTomato- (in blue) cells for each cell population.

**Supplemental figure 3: EMT tumor cells are present in breast cancer mouse models but lack CAF marker expression and function.** A) Example of gating strategy used to identify EpCAM-positive tumor cells (EpCAM+) and tumor cells that have lost EpCAM expression (EpCAM-). After exclusion of endothelial cells (CD31+) and immune cells (CD45+) remaining cells were divided into tdTomato-negative cells (tdTomato-, coming from transplanted tumor tissue) and tdTomato-positive cells (tdTomato+, coming from recipient mTmG host). EpCAM-positive tumor cells were defined as tdTomato-/EpCAM/E-Cadherin+/CD49f+/low. EpCAM-negative tumor cells were defined as tdTomato-/EpCAM/E-Cadherin-/CD49f-/PDGFRβ-. tdTomato+ cells consisted primarily of fibroblasts as they lacked expression of epithelial/basal markers (EpCAM, E-Cadherin, CD49f), but were positive for PDGFRβ. B) Quantification of EpCAM+ tumor cells and EpCAM- tumor cells in all MMEC transplanted mice (n=5 mice per time point). C) Collagen contraction assay with EpCAM+ tumor cells (n=2), EpCAM- tumor cells (n=2) and tdTomato+ CAFs (n=3) isolated from WB1P-Myc MMEC transplanted mTmG mice. Cells were plated at a density of 200.000 cells per collagen gel and left to contract for 72 hrs. D) Mammary tumor-free survival of FVB/n mice orthotopically injected with EpCAM+ tumor cells, EpCAM- tumor cells or tdTomato+ CAFs (n=2 mice per group). Statistical significance was determined by Log-rank Mantel Cox test. E) Tumor volumetric measurements of FVB/n mice orthotopically injected with EpCAM+ tumor cells, EpCAM-tumor cells or tdTomato+ CAFs (n=2 mice per group). Statistical significance compared to CAF-injected mice was determine by two-way ANOVA with Bonferonni's multiple comparison correction (EpCAM-1 p=0.0001, EpCAM-2 p=0.0070, EpCAM+1 p=0.0108, EpCAM+2 p=0.0001).

**Supplemental figure 4: Efficient bone marrow transplantation shows tdTomato+ BM-MSCs.** A) Gating strategy to define to and validate tdTomato expression in bone marrow mesenchymal stem cells (BM-MSCs). Bone marrow was harvested from FVB/n control mice, control mTmG mice and mTmG bone marrow-transplanted EPtn, EH1047R, B1P and B1P-Myc mice and stained with CD45-PerCP, Sca1-APC-Cy7 and CD90-FITC. MSCs were defined as CD45-/Sca1+/CD90high. B-E) Representative flow cytometry plots showing tdTomato expression in mTmG control mice (in red), FVB/n control mice (in blue) and two mTmG bone marrow transplanted mice (in orange and green) of the indicated tumor mouse models. Panel B shows EPtn, panel C shows EH1047R, panel D shows B1P and panel E shows B1P-Myc. F) Quantification of tdTomato- and tdTomato+ MSCs in bone marrow transplanted mice of the indicated tumor models (n=5 mice per model).

**Supplemental figure 5: Bone marrow transplantations in EPtn;mTmG mice confirm findings that BM-MSCs do not contribute to the population of CAFs in ILC.** A) Schematic representation of experimental set-up and its potential outcomes. B) Quantification of GFP expression within the mammary epithelium compartment (EpCAM+) in established tumors 18 weeks after lenti-cre injection (n=5 mice). C) Representative flow cytometry plot of tdTomato and GFP expression of EpCAM+ mammary epithelial cells of FVB/n control mice (control, in blue), control non-intraductal injected EPtn;mTmG mice (NI, in purple) and two EPtn;mTmG mice after lenti-cre intraductal injection (18 weeks post injection, tumor 1 and 2 in green and orange). D) Quantification of tdTomato expression in indicated cell population within tumors harvested from EPtn;mTmG mice 18 weeks after intraductal lenti-cre injection (n=5 mice). E) Representative flow cytometry plot of tdTomato expression of PDGFRβ+ fibroblasts of FVB/n control mice (control in blue), control non-intraductal injected EPtn;mTmG mice (NI, in purple) and two EPtn;mTmG mice after lenti-cre intraductal injection (18 weeks post injection, tumor 1 and 2 in green and orange).

**Supplemental figure 6: Morphology and cellular composition of whole mammary gland transplanted tissue resembles endogenous mammary glands.** A) Representative images of carmine stainings performed on endogenous and transplanted third whole mammary gland 12 weeks post-surgery. B) Quantification of cellular composition of endogenous and transplanted mammary glands as determined by flow cytometry (n=5 mice per group). Mammary epithelial cells were defined as EpCAM+/CD31-/CD45-/PDGFRβ-. Endothelial cells were defined as CD31+/CD49f+/EpCAM-/CD45-. Immune cells were defined as CD45+/CD31-/EpCAM-. Fibroblasts were defined as EpCAM-/CD45-/CD31-/PDGFRβ+. Statistical significance was determined using student's t-test.

**Supplemental figure 7: iCAFs and myCAF gene expression signatures are also present in human invasive ductal carcinoma (IDC).** A) iCAF and myCAF gene signatures identified in from single cell transcriptomics on WEPtn tumors and used throughout the manuscript as gene signatures that mark functionally distinct CAF subpopulations. B) iCAF gene expression signature in laser-microdissected human IDC samples (n=36) separated into tumor epithelium (marked in yellow) and tumor stroma (marked in purple). C) myCAF gene expression signature in human IDC. D) Single sample gene set enrichment analysis of iCAF and myCAFs scores in human IDC tumor epithelium and tumor stroma. E) UMAP plot of single-cell transcriptomics analysis of end-stage WB1P and WB1P-Myc tumors (n=2 mice per tumor type). Tumors were digested to single cell suspensions and sorted for live single cells by FACS (dapi-negative) and subjected to single-cell RNA sequencing using the Dropseq platform. First panel indicates the identified clusters and the second panel shows the distribution of cells from each mouse model. The clusters were annotated by marker gene expression. Myeloid cells express *Cd45*, *Cd14*, *Lyz2*, *Aif1* and various Cathepsins. Tumor cells express *Epcam*, *Krt8* and *Krt18*. Endothelial cells express *Pecam1*, *Cdh5* and *Esam*. CAFs express various collagens, *Tnc* and *Postn*. F) UMAP plots showing *Col1a1* and *Col1a2* expression used to identify the CAF cluster for further downstream analyses.

**Supplemental figure 8: PDAC iCAFs and myCAFs show significant overlap in gene expression and function with ILC iCAFs and myCAFs.** A) UMAP plots of PDAC iCAF, myCAF and apCAF gene signatures plotted on single cell dataset of NFs and CAFs isolated from EPtn mice. B) Gene signatures of PDAC iCAFs, myCAFs and apCAFs. C and D) Gene ontology analysis of the genes that define PDAC iCAFs (panel C) and myCAFs (panel D). E) Pearson's correlation between gene expression of PDAC iCAFs and ILC iCAFs beyond the gene signatures. F) Pearson's correlation between gene expression of PDAC myCAFs and ILC myCAFs beyond the gene signatures. In panel E and F the gene-signature defining genes are highlighted in orange (ILC), green (PDAC) or yellow (both). All other genes expressed by iCAFs and myCAFs are shown in grey.

**Supplemental figure 9: Gene ontology analysis of CD26- NFs and CD26+ NFs reveal activation of similar pathways but with different players.** A) Gene ontology analysis of the top-25 genes that define the cluster of CD26- NFs. B) Gene ontology analysis of the top-25 genes that define the cluster of CD26+ NFs. C) Genes expressed by CD26- and CD26+ NFs with the GO term extracellular matrix.

**Supplemental figure 10: Balance between CD26- and CD26+ fibroblasts shifts as tumors progress.** A) CD26- and CD26+ fibroblasts in WEPtn mice as determined by flow cytometry. ILC tumors were harvested at early (n=3), advanced (n=4) or end-stage (n=6) of tumor development. Together with mammary glands from control littermates (n=8) they were analyzed for CD26 expression. Fibroblasts were defined as EpCAM-/CD49f-/CD31-/CD45- cells. Within this population we determine the ratio of CD26- and CD26+ fibroblasts. B) CD26- and CD26+ fibroblasts in WEH1047R mice as determined by flow cytometry. Control mammary glands (n=8) and ILC tumors were harvested at early (n=4), advanced (n=5) or end-stage (n=6) of tumor development. C) CD26- and CD26+ fibroblasts in WB1P mice. Control mammary glands (n=11) and TNBC tumors were harvested at 3x3 mm (n=3), 8x8 mm (n=9) or 15x15 mm (n=7). D) CD26- and CD26+ fibroblasts in WB1P-Myc mice. Control mammary glands (n=8) and TNBC tumors were harvested at 3x3 mm (n=6), 8x8 mm (n=4) or 15x15 mm (n=5). A two-way ANOVA with Bonferonni multiple comparison correction was used to determine statistical significance was determined by comparing CD26- or CD26+ NFs to the CD26- or CD26+ CAFs of the different time points, respectively.

**Supplemental figure 11: Recruitment of CD26+ NFs does not dependent on CD26 enzymatic activity.** A) bright field image of FACS-sorted CD26- and CD26+ mammary NFs. Image taken one week after sorting. B) Transwell assay to assess the recruitment of CD26- and CD26+ NFs towards two WEPtn-derived ILC tumor cell lines in the absence and presence of CD26 inhibitors PK-44 (100 nM) and sitagliptin (100 nM). Representative result of two independent experiments with similar results. C) Transwell assay to assess the recruitment of CD26- NFs, CD26+ NFs or mixed CD26- and CD26+ NFs towards ILC tumor cells. Fibroblasts were harvested from mTmG mice by FACS and cultured for three days before switching the CD26+ NFs with Cre-encoding adenovirus to replace tdTomato expression with GFP. Four days post-switching the cells were plated in the recruitment assay. Representative result of four independent experiments with similar results. D) Quantification of transwell assays in panel C. Statistical significance was determined using student's t-test. E) Representative image of organotypic invasion assays performed with WB1P-derived tumor cells. F) Quantification of organotypic invasion assays based on EpCAM staining (n=3 independent experiments with similar outcome). Invasion was measured in um from top of the gel to the invasive front of the tumor cells. Statistical significance was determine using student's t-test.

**Supplemental figure 12: Differential response of CD26- and CD26+ NFs to tumor CM.** A) Transcriptomics analysis and hierarchical clustering of CD26- and CD26+ NFs cultured in control CM or WEPtn-derived tumor cell CM (n=6 WEPtn cell lines). Top 100 differential expressed genes are displayed here. B) KEGG pathway analysis of significantly upregulated (Log2FC>2 and FDR<0.05) genes between CD26- NFs cultured in control CM vs CD26- NFs cultured in tumor CM. C) KEGG pathway analysis of significantly upregulated (Log2FC>2 and FDR<0.05) genes between CD26+ NFs cultured in control CM vs CD26+ NFs cultured in tumor CM.

**Supplemental figure 13: MMEC transplantations in *En1-Cre;mTmG* mice show dynamics and differential gene expression in fibroblast populations during tumor development.** A) Analysis of fibroblasts distribution in control transplanted mice (n=5) and mice transplanted with pre-neoplastic WEPtn donor tissue (n=5). WEPtn-derived ILCs were analyzed at end-stage (42-weeks post transplantation). B) Analysis of fibroblasts distribution in control transplanted mice (n=5) and mice transplanted with pre-neoplastic WB1P donor tissue (n=5). WB1P-derived tumors were analyzed when they reached end-stage (15x15 mm). C) Analysis of fibroblasts distribution in control transplanted mice (n=5) and mice transplanted with pre-neoplastic WB1P-Myc donor tissue (n=5). WB1P-Myc-derived tumors were analyzed when they reached end-stage (15x15 mm). Statistical significance of panels A, B and C was determine by student's t-test comparing control to tumor for each of the populations. D) Transcriptomics analysis and hierarchical clustering of En1+, CD26+, double+ and double- fibroblasts isolated from control transplanted and WB1P or WB1P-Myc transplanted *En1-Cre;mTmG* mice. Top 100 genes are shown here. Fibroblasts were isolated by FACS and defined as EpCAM-/CD49f-/CD31-/CD45-.

**Supplemental figure 14: Differentially expressed cytokines between CD26- and CD26+ NFs do not impact recruitment of T- or B-cells.** A) Results of cytokine array incubated with CM derived from three-day-co-cultures of CD26- NFs with tumor cells and CD26+ NFs with tumor cells. Differentially expressed proteins are boxed. B) Quantification of entire cytokine array as shown in panel A. C and D) Results of transwell assays used to investigate recruitment of splenocytes towards the CM of fibroblast and tumor cell mono- or co-cultures. Migrated splenocytes were harvested from the bottom compartment after 24 hours, stained for CD3 and B220 to stain T- and B-cells, respectively and quantified by flow cytometry. Panel C shows migrated T-cells and panel D shows migrated B-cells. Results shown in panel C and D are from 3 independent experiments with similar outcome.
